# Control of Blood Pressure Variability Across Behavioral States by Brainstem Adrenergic Neurons

**DOI:** 10.1101/2025.05.14.654157

**Authors:** George M. P. R. Souza, Harsha Thakkalapally, Faye E. Berry, Ulrich M. Atongazi, Daniel S. Stornetta, Stephen B.G. Abbott

## Abstract

Short-term blood pressure (BP) variability is increasingly recognized as an independent predictor of cardiovascular and cerebrovascular risk, yet the central mechanisms that govern this variability, particularly across behavioral states, remain poorly defined. In this study, we investigated the role of C1 adrenergic neurons in the rostral ventrolateral medulla (RVLM^C1^) in the short-term BP regulation during sleep-wake transitions and physical activity in freely behaving rats. Using genetically targeted fiber photometry, we show that RVLM^C1^ neurons exhibit state-dependent activity, with rapid activation during arousal from non-REM sleep, sustained activity in REM sleep, and further recruitment during physical activity. We further demonstrate that baroreflex input is essential for the dynamic response of RVLM^C1^ neurons to pharmacological manipulations of BP and transitions to REM sleep. Strikingly, selective ablation of RVLM^C1^ neurons did not affect mean BP but caused pronounced instability during arousal and movement, underscoring their role in buffering BP fluctuations. These findings demonstrate that RVLM^C1^ neurons integrate arousal-related central drive with baroreceptor feedback to stabilize BP during changes in behavioral state. These findings suggest that the disruption of RVLM^C1^ neurons could underlie increased BP variability observed in pathological conditions, such as multiple system atrophy, even when mean BP is preserved.

## Introduction

Blood pressure is regulated within a narrow range to ensure adequate tissue perfusion and oxygen delivery in response to metabolic demands. While static BP measurements are an established clinical marker of cardiovascular health, accumulating evidence highlights increased short-term BP variability as an independent risk factor for cardiovascular and cerebrovascular events, as well as all-cause mortality–particularly in older adults and critically ill patients (Schutte et al., 2022; Sheikh et al., 2023; Kulkarni et al., 2025).

Short-term fluctuations in BP are a fundamental manifestation of arousal and physical activity, reflecting state-dependent modulation of the autonomic nervous system by central feed-forward control integrated with visceral sensory feedback from the cardiovascular system and somatic feedback from active muscles (Silvani et al., 2015; Dampney, 2016; Barman and Yates, 2017; Benarroch, 2018). Among these mechanisms, negative feedback from the arterial baroreflex is considered to be critical for short-term stability of BP through the regulation of central circuits confined to the medulla (Kumada et al., 1990; Guyenet, 2006; Dampney, 2017). While the arterial baroreflex is understood to be important in the stability of BP, the central mechanisms that integrate the baroreflex feedback with changes in behavioral states are incompletely defined.

The rostral ventrolateral medulla is an established node within the central autonomic network that contains barosensitive spinally-projecting excitatory neurons that regulate the activity of peripheral sympathetic vasoconstrictor nerves (Guyenet et al., 2018). A subpopulation of these neurons expressing catecholamine cell markers, referred to as C1 neurons (RVLM^C1^), are critical in the sympathetic response to physiological stressors (Guyenet et al., 2013), such as blood loss (Souza et al., 2022b) and hypoxia (Schreihofer and Guyenet, 2000; Madden et al., 2006; Wenker et al., 2017) as result of feedback from peripheral arterial baroreceptors and chemoreceptors (Brown and Guyenet, 1985; Koshiya et al., 1993; Sun and Reis, 1993; Chan and Sawchenko, 1994; Erickson and Millhorn, 1994). By contrast, the contribution of these cells to ‘tonic vasomotor tone’ at rest in undisturbed conditions is unsettled. Genetically-targeted optogenetic inhibition of RVLM^C1^ results neurons results in a modest fall in resting BP (<10 mmHg) while chronic ablation of RVLM^C1^ neurons results in either no change or a modest fall in resting mean BP (Schreihofer et al., 2000; Madden and Sved, 2003; Madden et al., 2006; Andrade et al., 2019; Toledo et al., 2019) despite significant deficits in several cardiovascular sympathoexcitatory reflexes (Schreihofer and Guyenet, 2000; Madden et al., 2006). The modest effect of the ablation of RVLM^C1^ neurons on resting MAP has led to a focus on the role of these neurons in the sympathetic response to physiological stressors and disease states (Guyenet et al., 2013, 2018). However, it remains to be determined if RVLM^C1^ neurons contribute to the homeostatic regulation of BP during changes in behavioral state.

Several lines of evidence suggest that RVLM^C1^ neurons may be recruited during periods of arousal and physical activity. They receive input from brain regions that regulate arousal and locomotor functions (Saper and Loewy, 1980; Van Bockstaele et al., 1991; Stornetta et al., 2016; Dempsey et al., 2017) and express receptors for arousal-related neuromodulators (Schwalbe et al., 2024), including orexin (Puskás et al., 2010). Furthermore, voluntary treadmill exercise markedly increases c-Fos expression in RVLM^C1^ neurons (Barna et al., 2012; Kumada et al., 2017), supporting a role for RVLM^C1^ neurons in the cardiovascular response to physical activity. On the other hand, air-puff stress and foot-shock in unanesthetized rats does not increase c-Fos expression in RVLM^C1^ neurons (Dayas et al., 2001; Carrive and Gorissen, 2008; Furlong et al., 2014) and cell-specific destruction of RVLM^C1^ neurons does not attenuate the BP response to conditioned fear or restraint stress (Vianna and Carrive, 2010), suggesting RVLM^C1^ neurons are not required for the cardiovascular response to psychological stress. As such, important questions regarding the role of RVLM^C1^ neurons in BP regulation in relation to behavioral state remain unanswered.

The goal of this study was to determine the contribution of RVLM^C1^ neurons to BP regulation during changes in behavioral state. The hypothesis was two-fold: first, that the activity of RVLM^C1^ neurons during distinct behavioral states would parallel that of sympathetic nerve activity established in previous studies (Miki et al., 2022); and second, that RVLM^C1^ neurons would be important for the BP response to states associated with increased arousal and during physical activity. To test this, we used genetically targeted *in vivo* fiber photometry to measure the activity of barosensitive RVLM^C1^ neurons during natural sleep-wake cycling, arousal from sleep evoked by salient stimuli, and physical activity in freely behaving rats. In light of studies indicating that variations in baroreflex sensitivity contribute to changes in BP during changes in sleep-wake state (Dampney, 2017; Miki et al., 2022), we performed sino-aortic denervation to establish the contribution of the arterial baroreflex to the response of RVLM^C1^ neurons to changes in sleep-wake state in unanesthetized conditions. Finally, we selectively eliminated RVLM^C1^ neurons using a genetically targeted approach to establish their role in BP regulation in relation to behavioral state. Our data reveal that the activity of barosensitive RVLM^C1^ neurons encodes distinct behavioral states in parallel with feedback from the arterial baroreflex and that they are critical for stabilizing BP during changes in behavioral state and physical activity.

## Results

### The activity of barosensitive RVLM^C1^ neurons are modulated by the sleep-wake state

To characterize the activity of RVLM^C1^ neurons, we unilaterally injected the RVLM of TH-Cre rats an AAV encoding Cre-dependent GCaMP7s followed by implantation of a fiber optic above the injection site and instrumentation for physiological recordings (**Fig. 1A, B, C**). Based on histology from 6 cases, native GCaMP7s expression was observed in 54 ± 2 RVLM neurons in a 1-in-3 series of coronal sections, with 78 ± 3% of these neurons expressing detectable TH^+^ (**Fig. 1D)**; 53 ± 4% of all TH^+^ neurons in the RVLM expressed GCaMP7s. Fiber optic placement was consistently located in the RVLM, typically within 300 μm of the caudal end of the facial nucleus (**Suppl. 1**).

**Figure 1.**
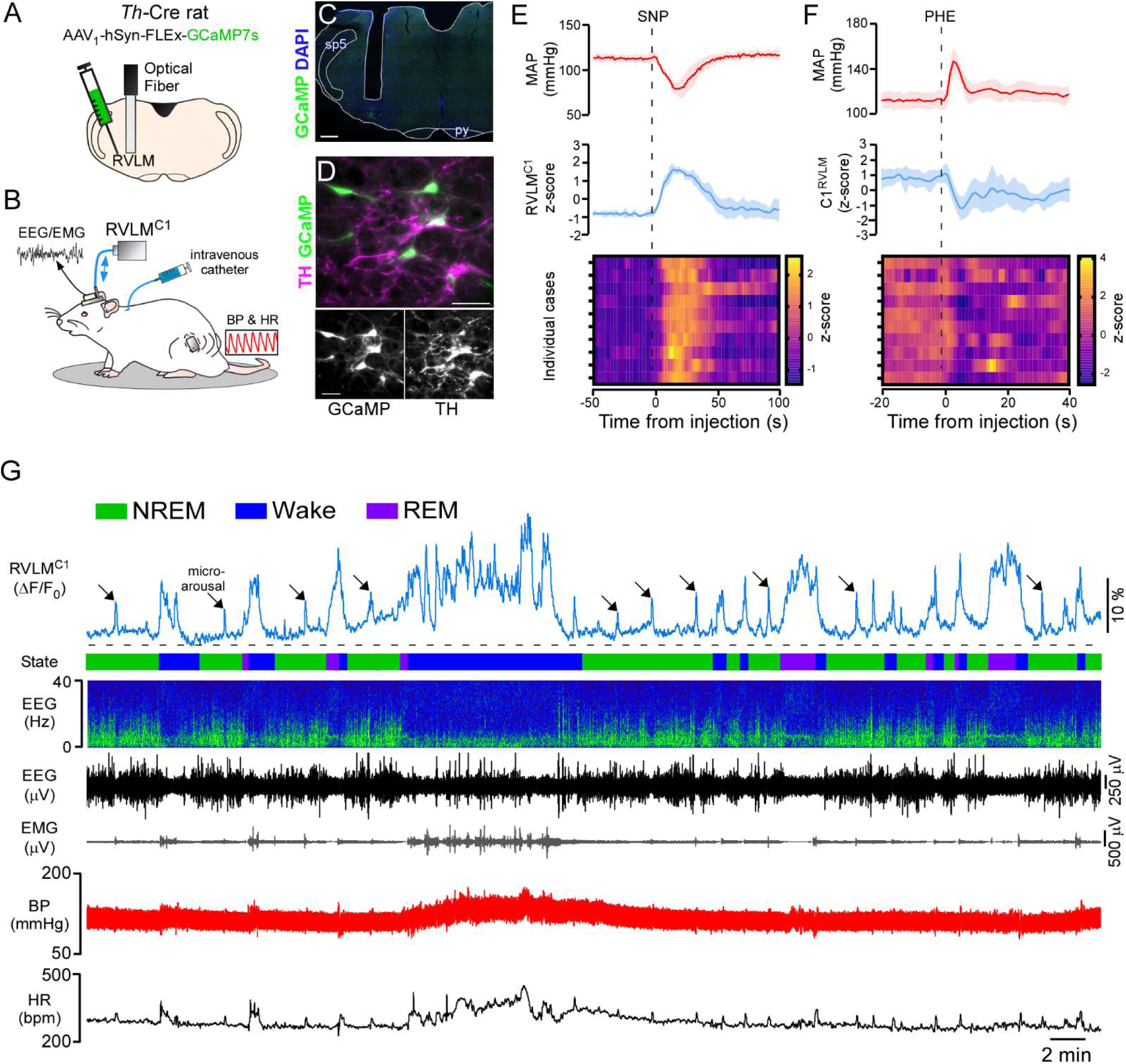
Genetically-targeted recordings of rostral ventrolateral medulla C1 neurons (RVLM^C1^) in freely behaving rats. **A.** Approach to transduce RVLM^C1^ neurons in *Th*-Cre rat for fiber photometry recordings using GCaMP7s. **B.** Experimental setup for fiber photometry recordings of RVLM^C1^ neurons with electrocorticography of brain activity (EEG), electromyography of neck muscle activity (EMG), and radiotelemetry recording of blood pressure (BP) and heart rate (HR) in an unanesthetized rat. **C-D.** Images showing expression of GCaMP7s in RVLM neurons expressing tyrosine-hydroxylase (TH). Scale bars: 400 µm for C, 100 µm for D **E.** Mean arterial blood pressure (MAP) and RVLM^C1^ fluorescence responses during hypotension following an intravenous bolus of sodium nitroprusside (SNP, 5 µg/kg in saline). Mean ± SEM **F.** MAP and RVLM^C1^ fluorescence responses during hypertension following an intravenous bolus of phenylephrine (PHE, 5 µg/kg in saline). Mean ± SEM **G.** Representative recording of RVLM^C1^ activity by fiber photometry simultaneously with sleep-wake state based on EEG and EMG, and BP and HR. Arrows indicate microarousals.

First, we characterized response of RVLM^C1^ neurons to baroreflex feedback to validate that our recordings reflect the activity of barosensitive neurons. Similar to our previous reports (Souza et al., 2022b), hypotension following an intravenous bolus of the vasorelaxant sodium nitroprusside (SNP) led to a robust increase in RVLM^C1^ activity (**Fig. 1E**), whereas hypertension following an intravenous bolus of the vasoconstrictor phenylephrine (PHE) led to a consistent reduction in fluorescence (**Fig. 1F)**. Next, we characterized RVLM^C1^ activity during natural sleep-wake cycling based on recordings of EEG and EMG (**Fig. 1G; Suppl. 2**). During NREM sleep,

RVLM^C1^ activity was stable and significantly lower than during wakefulness and REM sleep (**Fig. 1G, 2A, 2B**). We observed brief activations of RVLM^C1^ during microarousals in NREM sleep (**Fig. 1G; Suppl. 2A, B**). Transitions from NREM sleep to sustained wakefulness resulted in an abrupt increase in RVLM^C1^ that was coincident with the onset of EEG desynchronization (**Fig. 2A; Suppl. 2D**). During periods of extended wakefulness, RVLM^C1^ activity was elevated relative to NREM sleep, and in some cases presented bursts of activity during body movements associated with EMG activation (**Fig. 1G**). During transitions from wakefulness to NREM sleep, there was a progressive reduction in RVLM^C1^ activity, reaching a low-point at approximately the onset of slow-wave activity (**Fig 1G, 2A; Suppl. 2E**). During NREM-to-REM sleep transitions, RVLM^C1^ activity increased rapidly beginning at the onset of strong theta activity (**Fig 2A; Suppl. 2F**). Much like wakefulness, RVLM^C1^ activity during stable REM sleep exhibited spontaneous variations and rarely decreased to the levels of activity observed during NREM sleep (**Fig 1G**). During transitions from REM to wakefulness, RVLM^C1^ activity was variable, but often remained elevated at the time of the state transition and then decreased over approximately 10 s (**Fig 2A; Suppl. 2G**). Notably, the pattern of RVLM^C1^ activity closely paralleled the pattern of BP associated with NREM-to-wake and wake-to-NREM state transitions, whereas the pattern of RVLM^C1^ activity was orthogonal to changes in BP during NREM-to-REM and REM-to-wake transitions **(Fig. 2A**). The variations in RVLM^C1^ activity observed during sleep-wake state transitions was significantly different from recordings of GFP expressed in RVLM^C1^ neurons (**Fig. 2B**), indicating that these changes are not an artifact of blood flow or movement artifacts. Furthermore, the magnitude of the increase in RVLM^C1^ activity during NREM-to-wake and NREM-to-REM transitions was consistently smaller than the increase in activity during hypotension induced by SNP (**Fig. 2C**).

**Figure 2.**
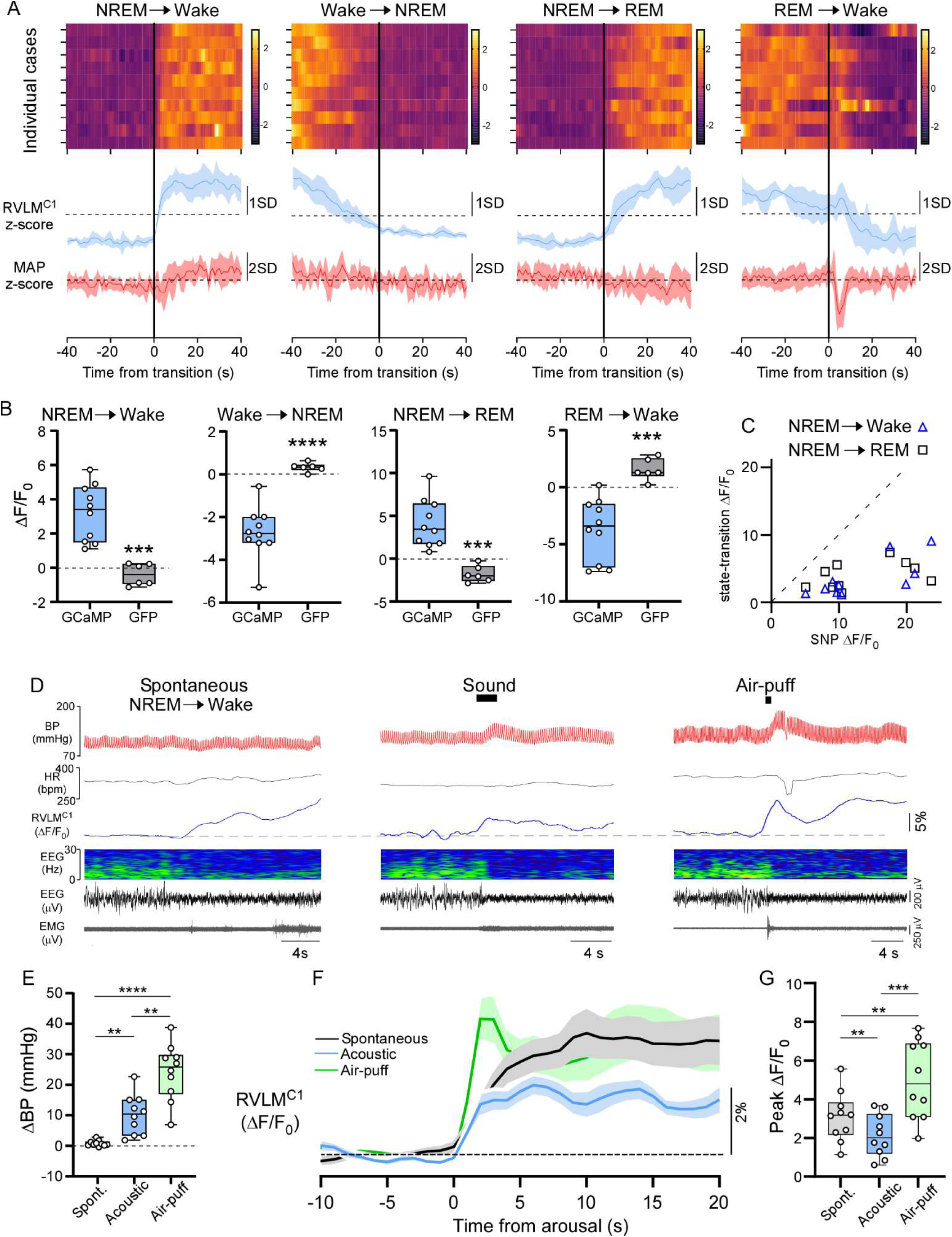
RVLM^C1^ activity is modulated by sleep-wake cycles and encodes arousal intensity. **A.** RVLM^C1^ activity during sleep-wake transitions. Upper panels depict z-score heat maps of ΔF/F_0_ during sleep-wake transitions for individual cases in grouped data (n=10). Middle panels depict grouped data of ΔF/F_0_ during sleep-wake transitions Lower panels depicts z-score of MAP during sleep-wake transitions. Mean ± SEM **B.** ΔF/F_0_ during state-transitions in GCAMP (n=10) and GFP (n=6) cases. For NREM-to-wake: Unpaired t-test, t=5.06, df=14, p=0.0002. For wake-to-NREM: Unpaired t-test, t=6.08, df=14 p<0.0001. For NREM-to-REM: Unpaired t-test, t=4.91, df=14, p=0.0002. For REM-to-wake: Unpaired t-test, t=4.53, df=14, p=0.0005. **C.** Scatter-plot of ΔF/F_0_ in response to hypotension induced with SNP against ΔF/F_0_ during NREM-to-wake and NREM-to-REM state transitions. **D.** Representative recordings of spontaneous NREM-to-wake transitions, and arousal from NREM induced by acoustic stimulation and air-puff. **E.** Peak ΔBP in response to conditions in panel D. (RM one-way ANOVA, F (1.697, 15.27) = 39.33, p<0.0001; asterisk reflect Tukey’s multiple comparisons test, ** P<0.01, **** P<0.0001. **F.** Mean ΔF/F_0_ (n=10) during spontaneous NREM-to-wake transitions, and arousal from NREM induced by acoustic stimulation and air-puff. Mean ± SEM **G.** Grouped data for peak ΔF/F_0_ during spontaneous NREM-to-wake transitions, and arousal from NREM induced by acoustic stimulation and air-puff (RM one-way ANOVA, interaction, F (1.262, 11.36) = 30.97, p<0.0001; asterisks reflect Tukey’s multiple comparisons test, ** P<0.01, *** P<0.001).

### RVLM^C1^ neurons encode the intensity of arousal

In humans, spontaneous and evoked EEG arousals are accompanied by graded autonomic activation resulting in transient increases in HR and BP (Davies et al., 1993; Sforza et al., 2000; Azarbarzin et al., 2014; Burke et al., 2020). To determine if the activation of RVLM^C1^ neurons during arousal from sleep varies with arousal intensity, we analyzed RVLM^C1^ fluorescence during microarousals (**Suppl. 2A-C**), spontaneous NREM-to-wake transitions, and arousal from NREM sleep induced by acoustic stimuli or air-puff (**Fig. 2F-I**). During microarousals, the magnitude of the HR response varies with the intensity of arousal. As such, we plotted the peak ΔHR against RVLM^C1^ fluorescence during microarousals and found a significant positive correlation between these variables despite inter-animal variability (**Suppl. 2C**). Moreover, there was a significant increase in the activation of RVLM^C1^ neurons during microarousals that were coupled with a sigh compared those that were not (microarousal without sigh vs. with sigh; 4.6 ± 0.7 vs. 5.3 ± 0.7, n= 10, Paired t test, t=4.438, df=9, P=0.0016). When arousal state transitions were induced by acoustic stimulation (**Fig. 2F**), we observed a significantly greater increase in BP compared to spontaneous NREM-to-wake transitions (**Fig. 2G**) although there was a less robust increase in RVLM^C1^ activity (**Fig. 2H, I**). On the other hand, air-puff stress resulted in a significantly larger increase in BP (**Fig 2G**) and a larger increase in RVLM^C1^ activity than both spontaneous NREM-to-wake transitions and acoustic stimulation (**Fig 2H, I**). Together, these results indicate that the activation of RVLM^C1^ during arousal from sleep encodes the intensity of arousal.

### Effect of baroreceptor denervation on the response of RVLM^C1^ neurons to perturbations in blood pressure

To establish the role of arterial baroreceptor feedback in the modulation of RVLM^C1^ activity in freely-behaving conditions, we performed sino-aortic denervation (Baro-X, n=8; **Fig. 3A**) or sham surgery (Sham, n=9). After 7 days recovery from sino-aortic denervation, MAP during NREM, REM or wakefulness was comparable to pre-surgical levels (**Suppl. 3A**), the cardiac baroreflex was antially reduced(**Fig. 3B-E)** and BP lability was elevated during wakefulness and REM sleep (**Suppl. 3B, C**), as seen in previous studies (Potts et al., 1997; Silveira et al., 2008; Waki et al., 2009; Abe et al., 2011; Lan et al., 2013; Amorim et al., 2016; Wenker et al., 2017), indicating that sino-aortic denervation was effective.

**Figure 3.**
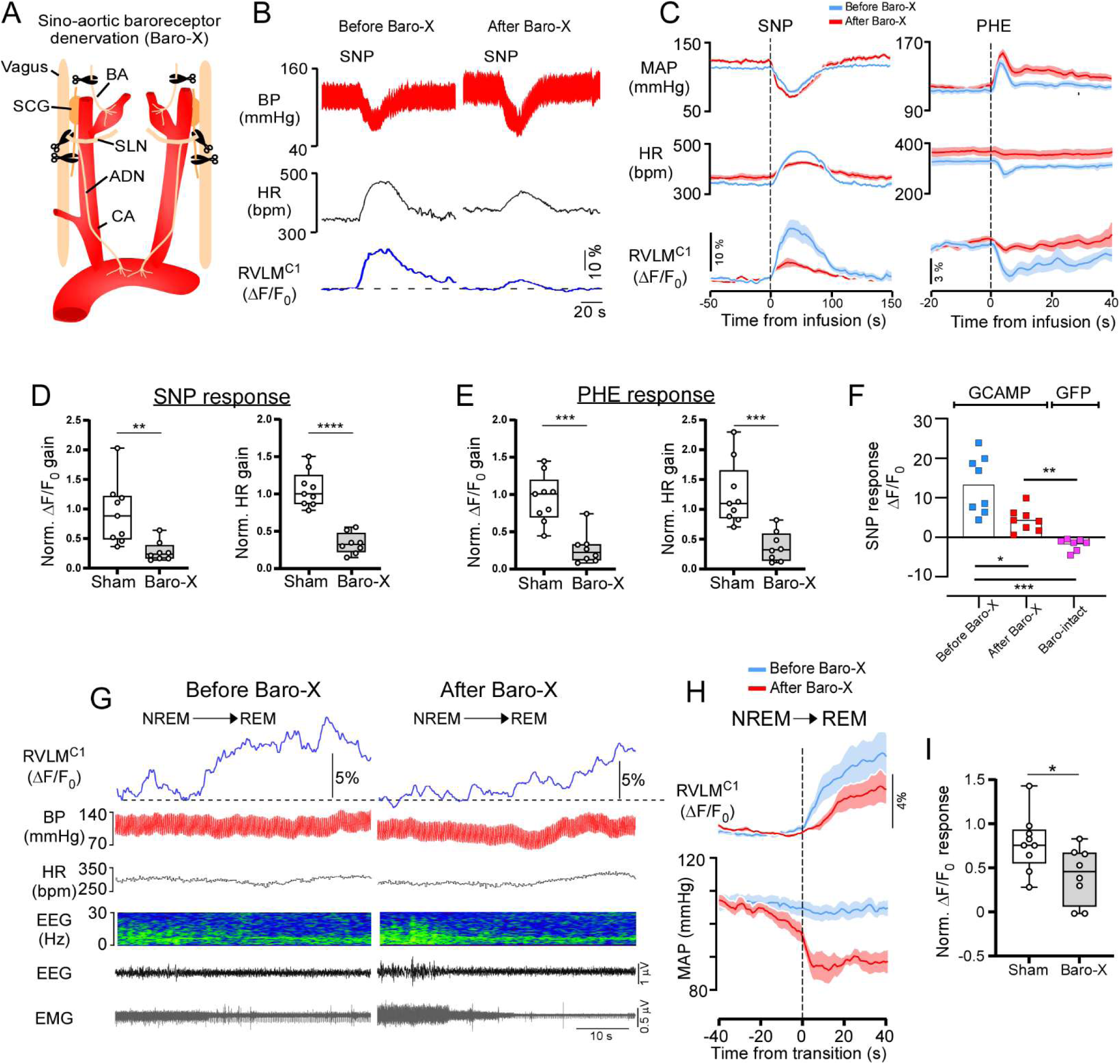
The arterial baroreflex is critical for the response of RVLM^C1^ neurons to perturbations in BP and transitions from NREM-to-REM. **A.** Illustration of the nerves cut during the sino-aortic denervation procedure. SCG: superior cervical ganglion, BA: xxx, SLN: xxx, ADN: xxx, CA: xxx **B.** Representative recordings of the BP, HR and RVLM^C1^ activity following an intravenous bolus of SNP in the same animal before and then 7 days after sino-aortic denervation (Baro-X). **C.** Time course of MAP, HR and ΔF/F_0_ in response to SNP and PHE before and after Baro-X (n=8). Mean ± SEM **D.** Grouped data for normalized ΔF/F_0_ gain and HR gain in response to SNP for sham (n=9) and Baro-X (n=8) rats. Normalized data reflects the ratio of post-surgery (ΔF/F_0_)/ΔmmHg to pre-surgery (ΔF/F_0_)/ΔmmHg so that a value of 1 reflects no change in the response after surgery. The same approach is used to normalize the HR response. ΔF/F_0_-Unpaired t-test, t=3.3, df=15, p=0.0049. HR-Unpaired t-test, t=7.5, df=15, p<0.0001. **E.** Grouped data for normalized ΔF/F_0_ gain and HR gain in response to PHE for sham (n=9) and Baro-X (n=8) rats. ΔF/F_0_-Unpaired t-test, t=4.9, df=15, p=0.0002. HR-Unpaired t-test, t=4.21, df=15, p<0.001. **F.** Grouped data for ΔF/F_0_ in response to SNP before and after Baro-X surgery in GCaMP expressing rats (n=9) and in Baro-intact rats expressing GFP (N=7) in RVLM^C1^ neurons. Brown-Forsythe ANOVA test, F (2.000, 9.870) =20.45, P=0.0003. Asterisk indicate results of Dunnett’s T3 multiple comparisons test. * P<0.05, ** P<0.01, P<0.001. **G.** Representative recordings of BP, HR and RVLM^C1^ activity, EEG and EMG during NREM-to-REM transitions before and after Baro-X in the same rat. **H.** Mean ΔF/F_0_ and MAP during NREM-to-REM transitions before and after Baro-X (n=8). Mean ± SEM **I.** Grouped data for normalized ΔF/F_0_ during NREM-to-REM transitions in Sham (n=9) or Baro-X (n=8) rats. Normalized data reflects the ratio of post-surgery to pre-surgery ΔF/F_0_ so that a value of 1 reflects no change in the response after surgery. Unpaired t test, t=2.3, df=15, p=0.035.

Consistent with arterial baroreceptor feedback mediating the response of RVLM^C1^ neurons to acute perturbations in BP, the fluorescent response of RVLM^C1^ neurons to both hypotension and hypertension was dramatically attenuated in the Baro-X groups compared to the Sham group (**Fig. 3C-E; Suppl. 3D-F**). Interestingly, despite the marked attenuation in the response of RVLM^C1^ neurons to hypotension after Baro-X, a small increase in fluorescence persisted that was significantly different from negative control recordings of RVLM^C1^ neurons expressing GFP in rats with intact baroreceptors (**Fig. 3F**; **Suppl. 3G**). This data demonstrates that feedback from the arterial baroreceptors is a critical source of dynamic negative feedback regulating RVLM^C1^ activity in relation to BP, but also that after chronic sino-aortic denervation these neurons retain some sensitivity to changes in BP or are activated by SNP through mechanisms that are unrelated to a fall in BP.

### Effect of sino-aortic denervation on the activity of RVLM^C1^ neurons during transitions behavioral state

Previous studies indicate that resetting of the arterial baroreflex contributes to the changes in autonomic function associated with sleep-wake patterns (Silvani et al., 2015; Miki et al., 2022). As such, we considered the possibility that the modulation of RVLM^C1^ neurons during sleep-wake state transitions may be disrupted by sino-aortic denervation. Interestingly, we observed an attenuation in the increased activity of RVLM^C1^ neurons during NREM-to-REM transitions in the Baro-X group compared the sham group (**Fig. 3G-I; Suppl. 4B, C**). Notably, NREM-to-REM transitions in Baro-X rats were also associated with a significant reduction in MAP (**Fig. 3H; Suppl. 4A**). On the other hand, RVLM^C1^ activity associated NREM-to-wake transitions (**Suppl. 4D, E**) and microarousals (**Suppl. 4F-I**) were comparable in Baro-X and Sham rats. Moreover, the recruitment of RVLM^C1^ activity during arousal from sleep induced by acoustic stimulation and air-puffstress was intact (**Suppl. 5A-D**). This result indicates that the activation of RVLM^C1^ neurons during NREM-to-REM transitions is mediated, in part, by modulation of arterial baroreflex feedback. However, the changes in RVLM^C1^ activity related to changes in sleep-wake state are mostly independent of changes in arterial baroreflex feedback. Collectively, these data indicate that the dynamic activity of RVLM^C1^ neurons involves distinct contributions from central sleep-wake and arousal-related inputs together with feedback from the arterial baroreceptors.

### Cell-selective ablation of RVLM^C1^ neurons increases short-term blood pressure variability

To establish the functional contribution of RVLM^C1^ neurons in the arousal state-dependent regulation of BP, we performed bilateral microinjections of the AAV encoding Cre-dependent taCasp3-TEVp to selectively induce cell apoptosis in RVLM^C1^ neurons (C1-Lesion), with controls rats injected with AAV encoding Cre-dependent GFP (C1-Intact) (**Suppl. 6A)**. Expression of Cre-dependent taCasp3-TEVp transgene resulted in a 92% reduction in PNMT-expressing neurons at bregma levels between −11.6 mm and −14.00 mm, and no change in the number of PNMT^+^ neurons in the C3 region, or Nissl^+^ neuronal profiles at the site of injections in the RVLM (**Fig. 4A, Suppl. 6B-D**).

**Figure 4.**
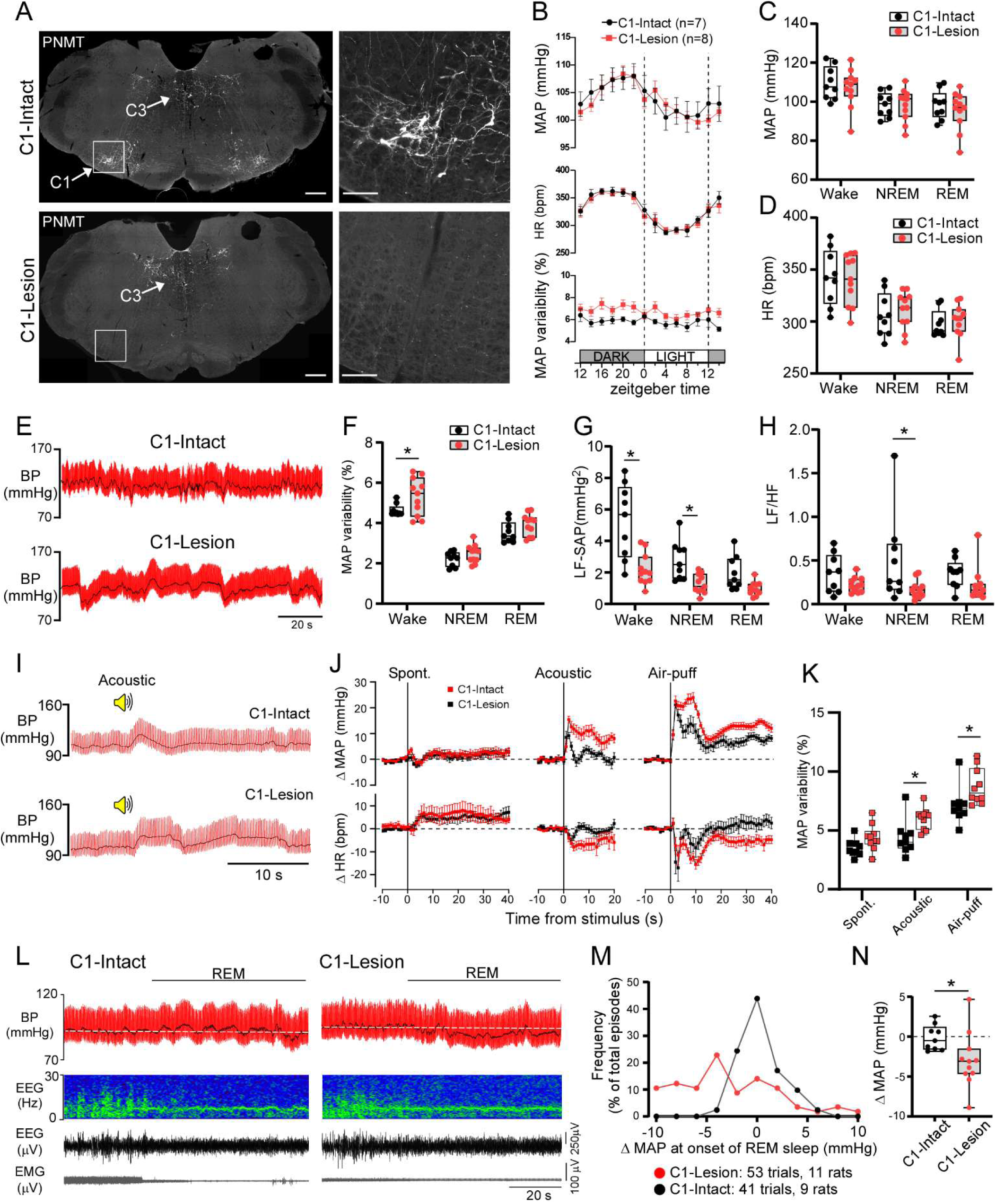
Genetically-targeted ablation of RVLM^C1^ neurons increases blood pressure variability. **A.** Image of the medulla from a C1-Intact and C1-Lesion case. Approximate bregma level: −12.3 mm. Scale bars: 500 µm in left panels and 100 µm in right panels. Mean ± SEM **B.** Grouped data for MAP, HR and the covariance of MAP (i.e., MAP variability) across dark and light periods in C1-Intact (n=7) and C1-Lesion (n=8) rats. MAP-RM two-way ANOVA, Time × Lesion, F _(13, 169)_ = 1.50, p=0.122; Time, F _(13, 169)_ = 19.04, p<0.0001; Lesion, F _(1, 13)_ = 0.032, p=0.86. HR-RM two-way ANOVA, Time × Lesion, F _(13, 169)_ = 0.2937, p=0.99; Time, F _(2.289, 29.75)_ = 20.04, p<0.0001; Lesion, F _(1, 13)_ = 0.18, p=0.68. MAP variability-RM two-way ANOVA, Time × Lesion, F _(13, 182)_ = 1.01, p=0.45; Time, F _(6.095, 85.33)_ = 2.007, p=0.073; Lesion, F _(1, 14)_ = 6.50, p=0.023. **C.** MAP during each state of sleep-wake cycle in C1-Intact (n=9) and C1-Lesion (n=11) rats. RM two-way ANOVA, State × Lesion, F _(2, 36)_ = 2.5, p=0.10; State, F _(1.9, 34)_ = 160, P<0.001; Lesion, F _(1, 18)_ = 0.26, p=0.62. **D.** HR during each state of sleep-wake cycle in C1-Intact (n=9) and C1-Lesion (n=11) rats. RM two-way ANOVA, State × Lesion, F _(2, 36)_ = 0.59, p=0.56; State, F _(1.5, 27)_ = 89, P<0.001; Lesion, F (1, 18) = 0.045, p=0.84. **E.** Representative recording of BP during a period of wakefulness in a C1-Intact and C1-Lesion rat. **F.** MAP variability across sleep-wake states in a C1-Intact (n=9) and C1-Lesion (n=11) rats. RM two-way ANOVA, State × Lesion, F _(2, 36)_ = 0.89, p=0.42; State, F _(1.6, 29)_ = 118, P<0.001; Lesion, F _(1, 18)_ = 7.4, p=0.014. Asterisk indicate results of Šídák’s multiple comparisons test-* p<0.05. **G.** Low frequency component of systolic arterial pressure (LF-SAP) across sleep-wake states in in C1-Intact (n=9) and C1-Lesion (n=11) rats. RM two-way ANOVA, State × Lesion, F _(2, 36)_ = 7.4, p=0.002; State, F _(1.1, 21)_ = 34, p<0.001; Lesion, F _(1, 18)_ = 18, p<0.001. Asterisk indicate results of Šídák’s multiple comparisons test-* p<0.05 **H.** HR variability sympathovagal balance (LF/HF) across sleep-wake states in C1-Intact (n=9) and C1-Lesion (n=11) rats. RM two-way ANOVA, State × Lesion, F _(2, 36)_ = 1.7, p=0.20; State, F (2, 36) = 0.36, p=0.70; Lesion, F _(1, 18)_ = 5.6, p=0.03. Asterisk indicate results of Šídák’s multiple comparisons test-* p<0.05. **I.** Representative recording of BP during acoustic stimulation in a C1-Intact and C1-Lesion rat. **J.** Grouped data for ΔMAP and ΔHR in response to spontaneous NREM-to-Wake transitions, and arousal from NREM induced by acoustic stimulation and air-puff in C1-Intact (n=9) and C1-Lesion (n=10) rats. Mean ± SEM. ΔMAP for spontaneous-RM two-way ANOVA, Time × Lesion, F _(59, 1003)_ = 0.36, p=0.99; Time, F _(5.8, 99)_ = 3.3, p=0.006; Lesion, F _(1, 17)_ = 0.35, p=0.56. ΔHR for spontaneous-RM two-way ANOVA, Time × Lesion, F _(59, 1003)_ = 0.54, p=0.99; Time, F _(2.6, 44)_ = 5.7, p=0.003; Lesion, F _(1, 17)_ = 0.39, p=0.54. ΔMAP for acoustic-RM two-way ANOVA, Time × Lesion, F _(40, 680)_ = 5.6, p<0.001; Time, F _(5.1, 86)_ = 14, p<0.001; Lesion, F _(1, 17)_ = 12, p=0.003. ΔHR for acoustic-RM two-way ANOVA, Time × Lesion, F _(40, 680)_ = 2.5, p<0.001; Time, F _(3.8, 64)_ = 3.6, p=0.012; Lesion, F _(1, 17)_ = 8.0, p=0.012. ΔMAP for air-puff-RM two-way ANOVA, Time × Lesion, F _(59, 1003)_ = 4.3, p<0.001; Time, F _(5.0, 86)_ = 58, p<0.001; Lesion, F _(1, 17)_ = 11, p=0.004. ΔHR for air-puff -RM two-way ANOVA, Time × Lesion, F _(59, 1003)_ = 2.1, p<0.001; Time, F _(59, 1003)_ = 6.5, p<0.001; Lesion, F _(1, 17)_ = 4.4, p=0.052. Mean ± SEM **K.** MAP variability following spontaneous NREM-to-wake transitions, and arousal from NREM induced by acoustic stimulation and air-puff in C1-Intact (n=9) and C1-Lesion (n=10) rats. RM two-way ANOVA, Stimulus × Lesion, F (2, 34) = 0.57, p=0.57; Stimulus, F (2, 34) = 57.97, P<0.001; Lesion, F (1, 17) = 13.6, p=0.002. Asterisk indicate results of Šídák’s multiple comparisons test-* p<0.05. **L.** Representative recording of BP, EEG, and EMG during a NREM-to-REM transition in a C1-Lesion rat. **M.** Frequency distribution of ΔMAP during NREM-to-REM transitions in C1-Intact and C1-Lesion cases. **N.** Grouped data for ΔMAP during NREM-to-REM transitions in C1-Intact (n=9) and C1-Lesion (n=11) cases. Unpaired t-test, t=2.246, df=18, p=0.038.

First, we examined a 24-hour recording of BP and HR by radiotelemetry in standard home-cage conditions with *ad libitum* food and water. In these conditions, there was no difference in MAP or HR between C1-Lesion and C1-Intact rats during the light and dark phase (**Fig. 4B, Suppl. 6E**). Moreover, the diurnal variation in MAP and HR across the dark and light phase was similar in C1-Lesion and C1-Intact rats (**Fig. 4B, Suppl. 6F**). On the other hand, we observed a significant increase in the coefficient of variation of MAP (i.e., MAP variability) throughout the 24-hour period (**Fig. 4B**), which suggests that RVLM^C1^ neurons contribute to the stability of BP but are redundant for the long-term regulation of BP and HR. To evaluate this observation in greater detail, we recorded BP concurrently with EEG/EMG in a plethysmography chamber to establish the role of RVLM^C1^ neurons in the cardiovascular changes associated with sleep-wake state. Similar to our 24-hour. recordings of BP, there was no difference in MAP or HR during NREM, REM sleep or wakefulness in C1-Lesion and C1-Intact rats (**Fig. 4C, D**). However, we found a significant increase in BP variability, primarily during wakefulness (**Fig. 4E, F**). To determine whether this effect reflects a disruption the sympathetic control of BP, we used spectral analysis of the systolic arterial pressure (SAP) variability and heart rate variability. We observed a marked reduction in the low-frequency component of SAP (LF-SAP), which reflects the sympathetic modulation of BP (Pagani et al., 1986), in C1-Lesion compared to C1-Intact rats during all sleep-wake stages, but most prominently during wakefulness (**Fig. 4G, Suppl. 6G**). Similarly, cardiac sympathovagal balance (LF/HF) was reduced in C1-Lesion rats owing to a reduction in the LF (**Fig. 4E; Suppl. 6H**), which reflects the sympathetic modulation of heart rate (Malliani et al., 1991). The increase in MAP variability and reduction in LF-SAP following ablation of RVLM^C1^ neurons were observed in both male and female rats (**Suppl. Table 1**). This data suggests that the loss of RVLM^C1^ neurons chronically disrupts the sympathetic regulation of vasomotor tone and HR resulting in an increase in BP variability.

We next examined whether the ablation of RVLM^C1^ neurons leads to dysfunctional regulation of BP during conditions in which they are most active, specifically, during transitions from NREM-to-wake and NREM-to-REM transitions, and in response acoustic stimulation and air-puff. While the cardiovascular response associated with spontaneous NREM-to-wake transitions was similar between groups (**Fig. 4J**), we observed a dramatically enhanced BP response following arousal evoked by acoustic stimulation and air-puff in C1-Lesion rats (**Fig. 4I, J**). Moreover, there was significant increase in the variability of BP following acoustic stimulation and air-puff (**Fig. 4K**). These data indicate that RVLM^C1^ neurons buffer against elevations in BP during abrupt arousals from sleep. Furthermore, we observed a marked increase in the probability of hypotensive events during NREM-to-REM transitions in C1-Lesion rats (**Fig. 4L-N**). Importantly, there was no difference between C1-Lesion and C1-Intact rats in the pattern of HR during NREM-to-REM transitions (**Suppl. 4K, L)** indicating that hypotensive events during these transitions are likely to be caused a dysregulation of sympathetic vasomotor tone. Together, our data shows that while RVLM^C1^ neurons are not essential for the long-term regulation of BP and HR, regardless of sleep-wake state, and are not required for cardiovascular stimulation during arousal-inducing stimuli, they are critical for the brain’s ability to defend BP against acute periods of hypertension and hypotension during changes in arousal state.

### A loss of RVLM^C1^ neurons leads to hypotension during rearing and jumping

Our data shows BP variability increases in C1-Lesion rats most prominently during wakefulness, which we hypothesized was related to inadequate buffering during periods of physical activity and movement. To test this hypothesis, we determined the role of RVLM^C1^ neurons in the regulation of BP during rearing, a stereotype postural change commonly observed in rats during exploration of a novel environment (Poulter et al., 2018). First, we used fiber photometry to determine whether RVLM^C1^neurons are activated during rearing in a large arena promoting natural exploration and observed a robust increase in their activity coinciding with rearing onset (**Fig. 5A, B**). Next, we examined the cardiovascular effects of rearing in C1-Lesion and C1-Intact rats, observing a negligible change in both MAP and HR in C1-Intact rats and an abrupt reduction in BP coupled with a tendency for an enhanced HR response in C1-Lesion rats (**Fig. 5C, D, G, H**). We also characterized the cardiovascular response to jumping in C1-Lesion and C1-Intact rats by adding a partition to the arena that they crossed during exploration (**Fig. 5E**). Similar to rearing, we observed a negligible change in both MAP and HR in C1-Intact rats during jumping and an abrupt reduction in BP coupled with an enhanced HR response in C1-Lesion rats (**Fig. 5E, G, H**). On the other hand, we observed no differences in BP and HR patterns during grooming between groups (**Fig. 5F, G, H**). Collectively, these data indicate that RVLM^C1^ neurons are critical in the prevention of hypotensive periods during postural changes and physical activity.

**Figure 5.**
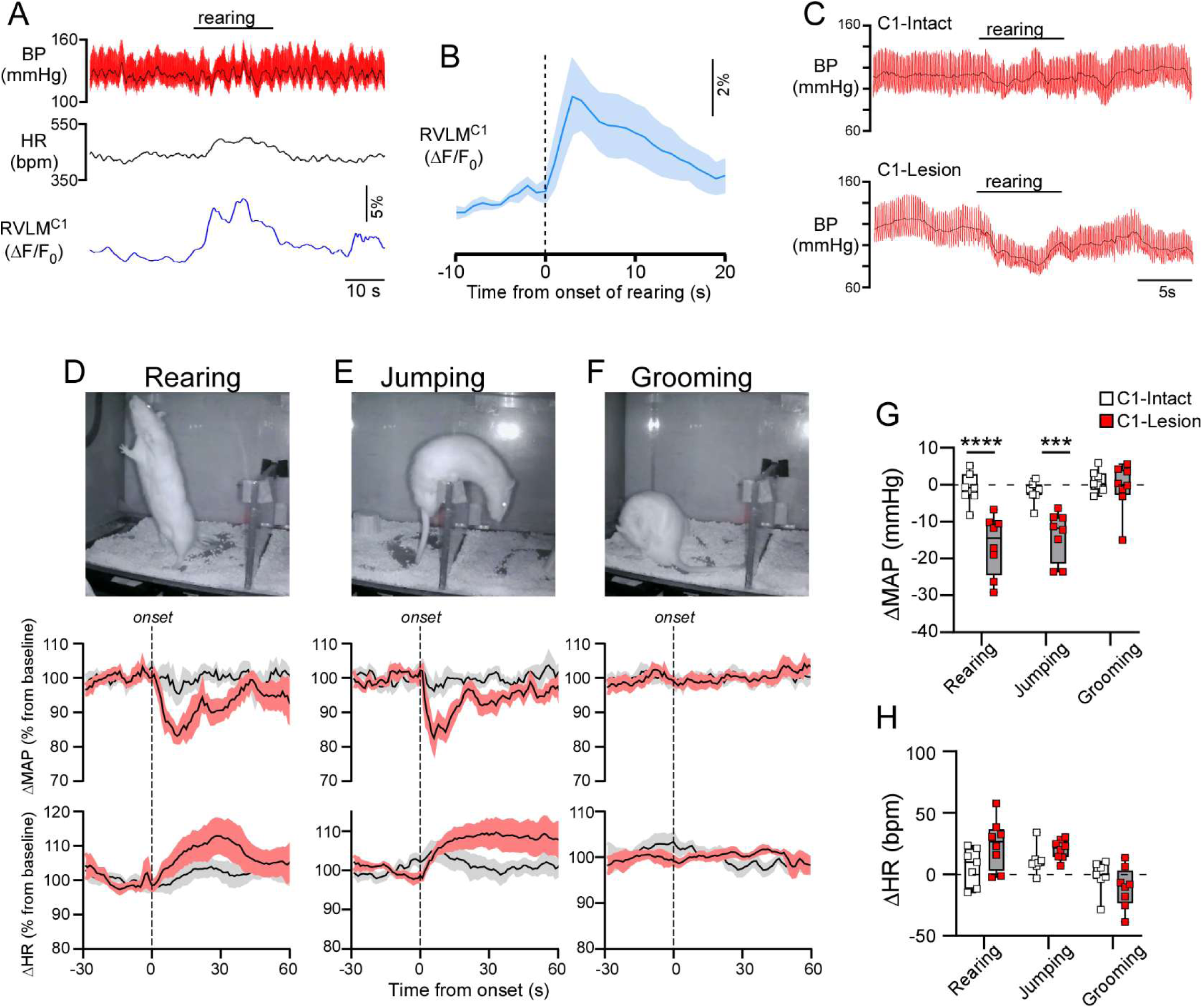
Ablation of RVLM^C1^ neurons leads to hypotension during rearing and jumping. **A.** Representative recording of BP, HR and RVLM^C1^ activity during rearing. **B.** Grouped data for ΔF/F_0_ during rearing (n=6). **C.** Representative recording of BP and HR during rearing in a C1-Intact and C1-Lesion rat. **D-F.** ΔMAP and ΔHR during rearing (D), jumping (E), and grooming (F) in a C1-Intact (n=6) and C1-Lesion (n=5) rats. Top images represent video recordings used to identify behavior. **G.** ΔMAP during rearing, jumping and grooming in C1-Intact (n=7) and C1-Lesion (n=8) rats. Mean ± SEM. Two-way ANOVA, Activity × Lesion, F _(2, 39)_ = 5.6, p=0.007; Activity, F _(2, 39)_ = 11, p<0.001; Lesion, F _(1, 39)_ = 31, p<0.001. Asterisk indicate results of Šídák’s multiple comparisons test-*** p<0.001, **** p<0.0001 **H.** ΔHR during rearing, jumping and grooming in C1-Intact (n=7) and C1-Lesion (n=8) rats. Two-way ANOVA, Activity × Lesion, F _(2, 39)_ = 3.3, p=0.047; Activity, F _(2, 39)_ = 11, p<0.001; Lesion, F _(1, 39)_ = 1.7, p=0.20.

## Discussion

This study shows that the activity of RVLM^C1^ neurons reflects the convergence of inputs regulated by behavioral-state and the arterial baroreflex, and that these neurons play a critical role in the stabilization of BP across behavioral states. We also demonstrate that the response of RVLM^C1^ neurons to acute changes in BP is dependent almost entirely on feedback from ‘high-pressure’ arterial baroreceptors, and that baroreflex feedback contributes to the increase in RVLM^C1^ activity during the onset of REM sleep, but not other sleep-wake states.

### RVLM^C1^ neurons encode arousal intensity and behavioral state

Our data demonstrates that the activity of RVLM^C1^ neurons is temporally coupled with sleep-wake state, displaying elevated activity in response to arousal from NREM sleep and sustained elevated activity during wakefulness and REM sleep. With the exception of wake-to-NREM transitions, which were characterized by a gradual reduction in RVLM^C1^ activity preceding the onset of stable slow-wave activity, the recruitment of RVLM^C1^ neurons in response to sleep-wake transitions was abrupt and coincided with changes in cortical activity. We also show that RVLM^C1^ activity correlates with the intensity of autonomic activation associated with arousal from sleep. The pattern of RVLM^C1^ activity during sleep-wake transitions also correlates with the progressive change in brain state and autonomic function that precedes sleep onset (Ogilvie, 2001; Shinar et al., 2006) and the abrupt desynchronization of cortical activity and recruitment of the peripheral sympathetic efferents with arousal from sleep (Floras et al., 1978; Horner, 1996). Of note, RVLM^C1^ neurons were consistently activated during microarousals and arousals from NREM sleep but do not appear to be required for the increase in BP or HR that characterizes these events, even when arousal is evoked by salient stimuli. As such, the prominent sympathetic component mediating the increase in BP during arousal from sleep (Khatri and Freis, 1967; Horner et al., 1995; Lo Martire et al., 2018) must rely on the activation of other pre-sympathetic neurons, for example, neurons in the medullary raphe (Zaretsky et al., 2003; Cao et al., 2004; Horiuchi et al., 2004).

REM sleep is a unique physiological state associated with regionally-specific changes in sympathetic activity resulting in mesenteric and renal vasodilation and muscle vasoconstriction (Mancia et al., 1971; Hornyak et al., 1991; Okada et al., 1991; Somers et al., 1993; Miki et al., 2004; Nagura et al., 2004). While neurons in regions within and adjacent to the RVLM are activated during REM sleep (Luppi and Fort, 2019; Cleary et al., 2025), there is uncertainty regarding the response of RVLM^C1^ neurons in this condition (Léger et al., 2009; Stettner et al., 2013; Yao et al., 2022). Our data demonstrate that RVLM^C1^ neurons, at a population-level, are activated during REM sleep. Interestingly, sino-aortic denervation attenuated the increase in RVLM^C1^ activity during the onset of REM sleep, which contrasts with RVLM^C1^ activity during NREM-to-wake transitions. This indicates that arterial baroreflex unloading or modulation of baroreflex sensitivity disinhibits RVLM^C1^ neurons during transitions to REM sleep, whereas the response of RVLM^C1^ neurons during other sleep-wake transitions is likely due to input from the central network regulating sleep-wake patterns. Sino-aortic denervation also led to pronounced hypotension at the onset of REM sleep, similar to previous work (Silveira et al., 2008), and this effect was partly replicated by selective ablation of RVLM^C1^ neurons. This result emphasizes the importance of the baroreflex-mediated disinhibition of RVLM^C1^ neurons as an important mechanism buffering against a fall in BP during transitions to REM sleep.

Notably, the increase in RVLM^C1^ activity observed during REM sleep was less pronounced than that evoked by baroreflex unloading with SNP, which we interpret as reflecting near-maximal recruitment of the RVLM^C1^ neuronal population. This disparity may reflect functional heterogeneity among RVLM^C1^ neurons in their responsiveness to REM-related inputs that cannot be resolved by bulk calcium imaging. Alternatively, the synaptic drive to RVLM^C1^ neurons during REM sleep may be inherently weaker or less synchronized than the robust activation induced by baroreceptor deafferentation. It is well-established that REM sleep increases sympathetic nerve activity targeting skeletal muscle (Hornyak et al., 1991; Somers et al., 1993; Miki et al., 2004). However, RVLM^C1^ neurons are understood to regulate vasoconstrictor activity in the splanchnic, renal and lumbar sympathetic nerves (Abbott et al., 2009; Kanbar et al., 2010; Mueller et al., 2011; Souza et al., 2022b). This evidence can be reconciled if the increase in RVLM^C1^ activity during REM sleep reflects an increase in the activity of a sub-population of neurons that selectively regulate muscle sympathetic nerve activity, or alternatively, if the reduction in splanchnic and renal sympathetic nerve activity during REM sleep is related to a reduction in the excitability of sympathetic preganglionic neurons in the spinal cord that preferentially regulate blood flow to these organs, perhaps owing to silencing of A5 noradrenergic (Souza et al., 2022a), which are inhibited during REM sleep (Fenik et al., 2002; Rukhadze et al., 2008; Saper et al., 2010).

*Role of the arterial baroreceptors in modulating RVLM^C1^ activity*.

The notion that RVLM^C1^ neurons respond to systemic blood pressure changes via baroreflex-mediated negative feedback is axiomatic (Guyenet, 2006; Barman and Yates, 2017; Dampney, 2017; Benarroch, 2018). However, direct evidence for the necessity of sino-aortic baroreceptors for the barosensitivity of RVLM^C1^ neurons is limited (Brown and Guyenet, 1985; Potts et al., 1997; Lan et al., 2013). Our data, based on genetically targeted fiber photometry recordings in freely behaving rats, shows that the dynamic response of RVLM^C1^ to both transient increases and decreases in BP is greatly reduced after sino-aortic denervation. Importantly, the fluorescent response to both hypotension and hypertension was unaffected in Sham-operated rats demonstrating that variations in recording quality over repeated recording sessions does not account for insensitivity of RVLM^C1^ neurons to BP perturbations after sino-aortic denervation. Thus, our results clearly indicate that RVLM^C1^ activity is barosensitive predominately due to inputs from the sino-aortic baroreceptors.

Interestingly, there was a still significant increase in RVLM^C1^ fluorescence during hypotension after sino-aortic denervation relative to baro-intact GCAMP-negative control recordings. As our approach for sino-aortic denervation replicated the cardiovascular phenotype reported in previous studies, it seems unlikely that incomplete denervation could account for this result. While we titrated the dose of SNP to avoid severe hypotension in sino-aortic denervated rats, it is plausible that subsidiary peripheral baroreceptor mechanisms are engaged after sino-aortic denervation, such as cardio-pulmonary receptors (Guo et al., 1982) and renal baroreceptors (Recordati et al., 1978; Schreihofer et al., 1997). Another possibility is that central baroreceptor or oxygen-sensitive mechanisms in glial cells (Gourine and Funk, 2017; Marina et al., 2020) or RVLM^C1^ themselves (Sun and Reis, 1994) are recruited due to changes in cerebral blood flow and oxygenation produced by acute hypotension. However, in unanesthetized goats, intravenous infusion of SNP at doses similar to that used in this study did not affect cerebral blood flow (Ivankovich et al., 1976). As such, further study will be required to parse the inputs that stimulate RVLM^C1^ neurons during hypotension in the absence of arterial baroreflex feedback.

### The loss of RVLM^C1^ neurons increases blood pressure variability

Prior studies employing saporin-based lesions to ablate RVLM^C1^ neurons in rats have yielded variable effects on resting mean BP (Schreihofer et al., 2000; Madden and Sved, 2003; Madden et al., 2006; Andrade et al., 2019; Toledo et al., 2019). However, this method indiscriminately targets other medullary catecholaminergic populations, including A5 neurons, due to retrograde uptake of the toxin by noradrenergic axons (Schreihofer and Guyenet, 2000; Madden et al., 2006). As a result, the specific contribution of RVLM^C1^ neurons to baseline BP regulation remains unresolved in these studies. The present study demonstrates that genetically targeted ablation of RVLM^C1^ neurons does not reduce resting meaning BP across distinct behavioral states or alter the diurnal BP variations despite a marked reduction in the sympathetic component of SBP and HR variability. A simple explanation for these results is that compensatory adjustments in blood volume and vascular tone mediated by renal and endocrine mechanisms (Hall et al., 2012) restore mean BP after a reduction in the sympathetic modulation of BP due to RVLM^C1^ cell loss. Interestingly, the ablation of RVLM^C1^ neurons markedly increased BP variability, particularly following arousal from sleep evoked by salient stimuli and during physical activity, which were conditions in which RVLM^C1^ neurons were most active based on our fiber photometry recordings. These results highlight the critical and non-redundant role of RVLM^C1^ neurons in minimizing BP variability during changes in behavioral state and physical activity.

BP regulation during changes in behavioral state and physical activity involves both central command and feedback regulation of the cardiovagal and sympathetic drive. The arterial baroreflex is considered a critical input regulating BP in these conditions (Dampney, 2017), and it is likely that the disruption of the sympathetic baroreflex contributes to the marked increase in BP variability after ablation of RVLM^C1^ neurons. However, the role of baroreflex feedback in BP regulation during rearing is equivocal (Waki et al., 2009; Abe et al., 2011), highlighting that the increase in BP variability after ablation of RVLM^C1^ neurons is probably also due to the disruption of central command mechanisms (Mori et al., 2005; Waki et al., 2009; Abe et al., 2011; Sugiyama et al., 2011). Supporting this, our photometry recordings show that the increase in RVLM^C1^ activity during rearing coincided with movement onset. Feed-forward control of RVLM^C1^ neurons could arise from the vestibular system (Barman and Yates, 2017), hypothalamus (Mileykovskiy et al., 2005; Puskás et al., 2010), and coupling between RVLM^C1^ neurons and circuits governing motor control (Arber and Costa, 2022; Koba et al., 2022; Noga and Whelan, 2022). Of note, the vestibular system contributes to the cardiovascular response to rearing in rats (Yates et al., 2014), and there is electrophysiological data (Miller et al., 2020) and well-established circuitry (Holstein et al., 2011; Barman and Yates, 2017) to account for the vestibular modulation of RVLM^C1^ neurons independent of the baroreflex. In sum, the pronounced increase in BP variability during physical activity after cell-selective ablation of RVLM^C1^ neurons highlights the pivotal role of these cells in fine-tuning sympathetic drive to the vasculature to buffer changes in BP during daily activities.

## Conclusion

In conclusion, this study indicates that the dynamic activity of RVLM^C1^ neurons stabilizes BP across behavioral states by integrating central command and baroreflex feedback. Notably, while mean arterial pressure reflects the integrated output of multiple redundant regulatory systems, moment-to-moment BP stability appears to be uniquely dependent on RVLMC1 neuron activity as a central node of control. Thus, a disruption in the function of these neurons could account for pathological BP variability in humans with normal mean BP. Indeed, previous work demonstrates that there is a profound loss of RVLM^C1^ neurons in multiple system atrophy (Benarroch et al., 1998; Coon et al., 2016), a form of Parkinson’s disease with a high incidence of orthostatic hypotension. As such, further exploration of the role of RVLM^C1^ neurons in the regulation BP will lead to a better understanding of etiology of in short-term BP variability in human disease (Sheikh et al., 2023; Kulkarni et al., 2025).

## Supplementary Figure Legends

**Supplementary figure 1.**
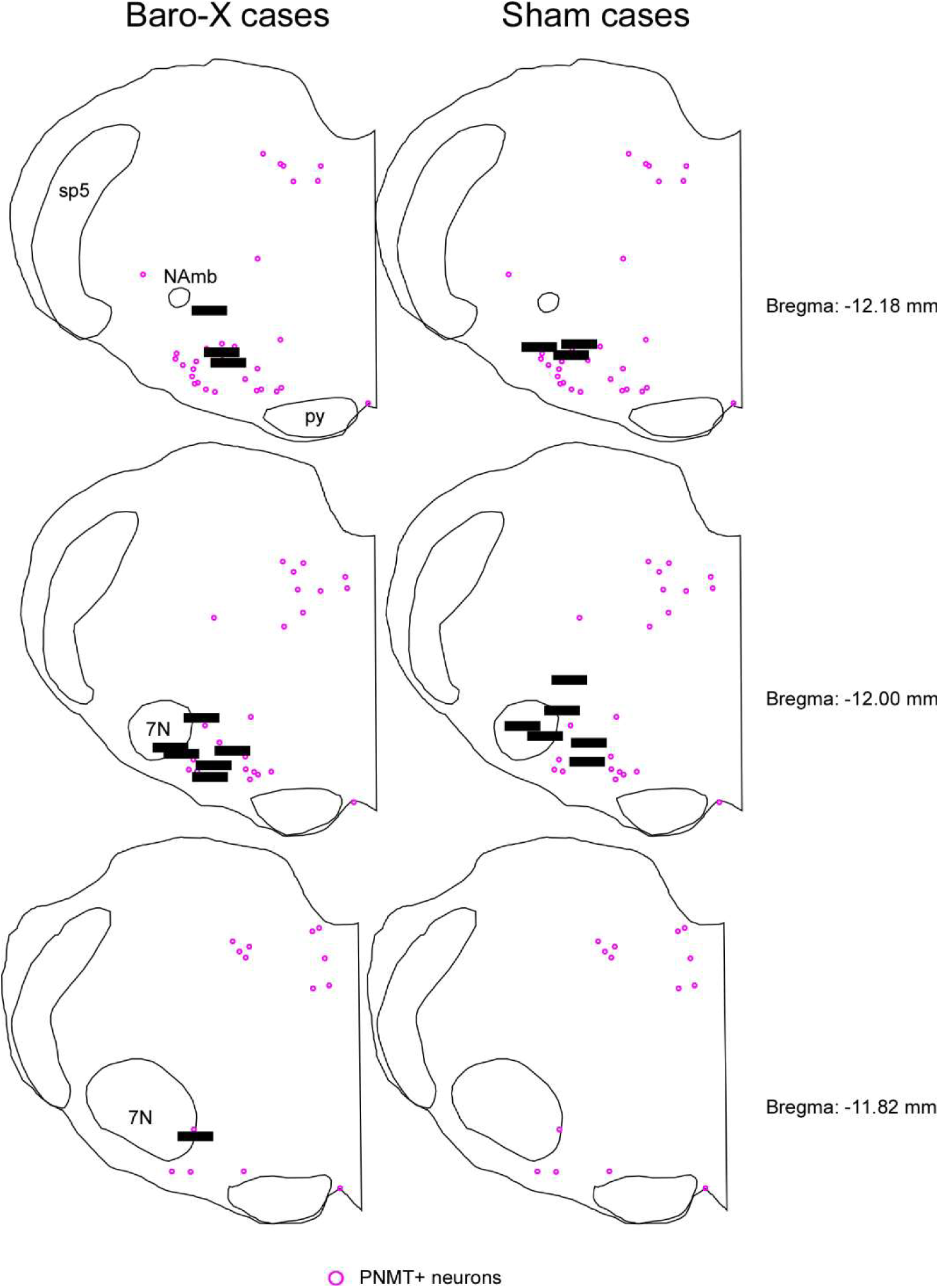
Summary of fiber placement for GCAMP used in analysis separated by whether they were included in the Baro-X or Sham group. The distribution of C1 neurons, identified by PNMT immunofluorescence, is indicated by magenta circles. Abbreviations: NAmb - compact formation of the nucleus ambiguus, sp5 - spinal trigeminal tract, py - pyramidal tract, 7N - facial motor nucleus.

**Supplementary figure 2.**
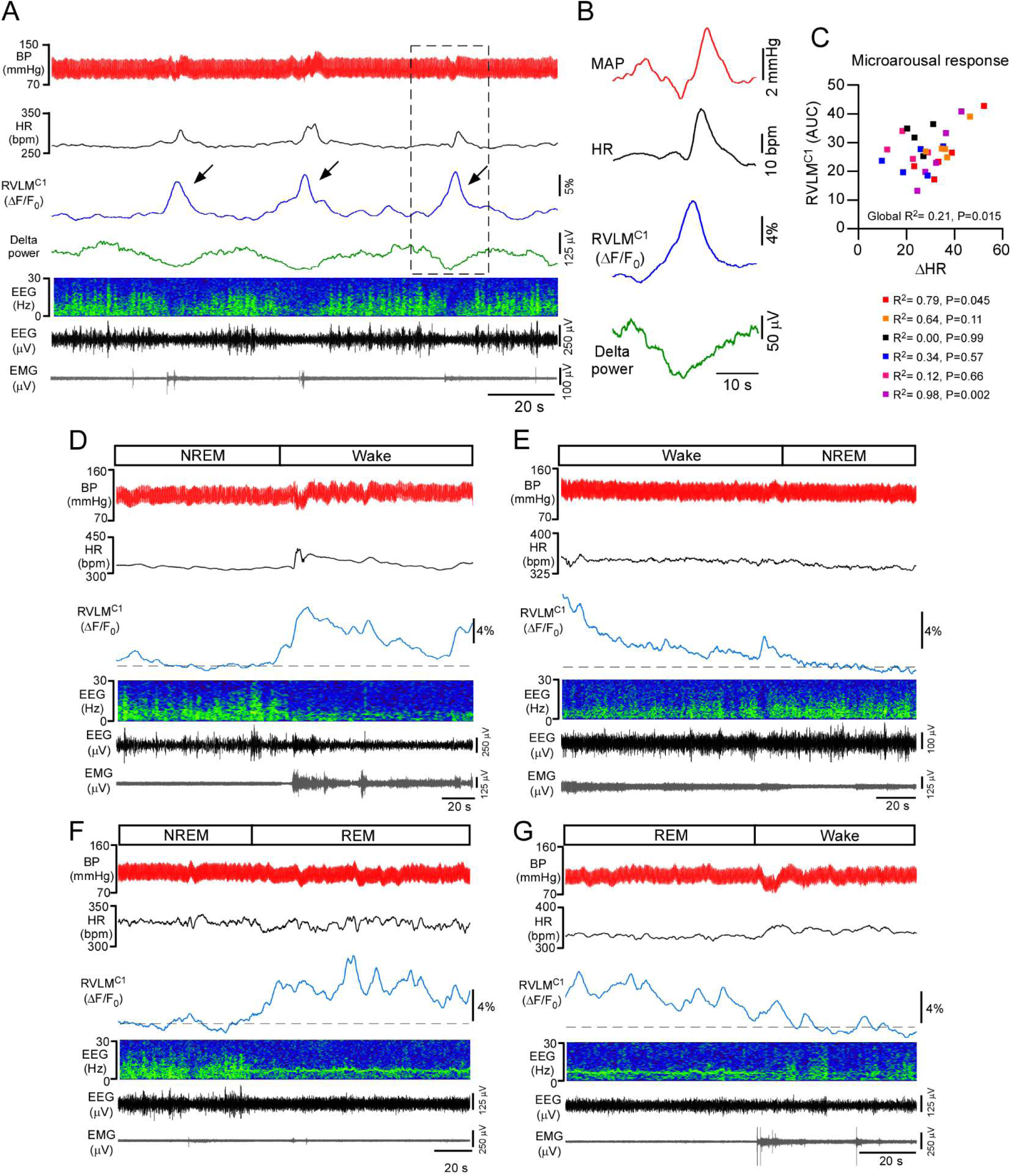
Representative recordings of BP, HR, RVLM^C1^ activity (ΔF/F_0_), EEG, and EMG during microarousals and sleep-wake state transitions. **A)** Example of microarousals indicated by arrows. **B)** Expanded exampled indicated in A showing pattern of RVLM^C1^ activity relative to other physiological parameters. **C)** Scatter-plot of peak ΔHR and peak ΔF/F_0_ following microarousals for 6 cases. **D)** Example of NREM-to-wake transition **E)** Example of wake-to-NREM transition **F)** Example of NREM-to-REM transition **G)** Example of REM-to-wake transition

**Supplementary figure 3.**
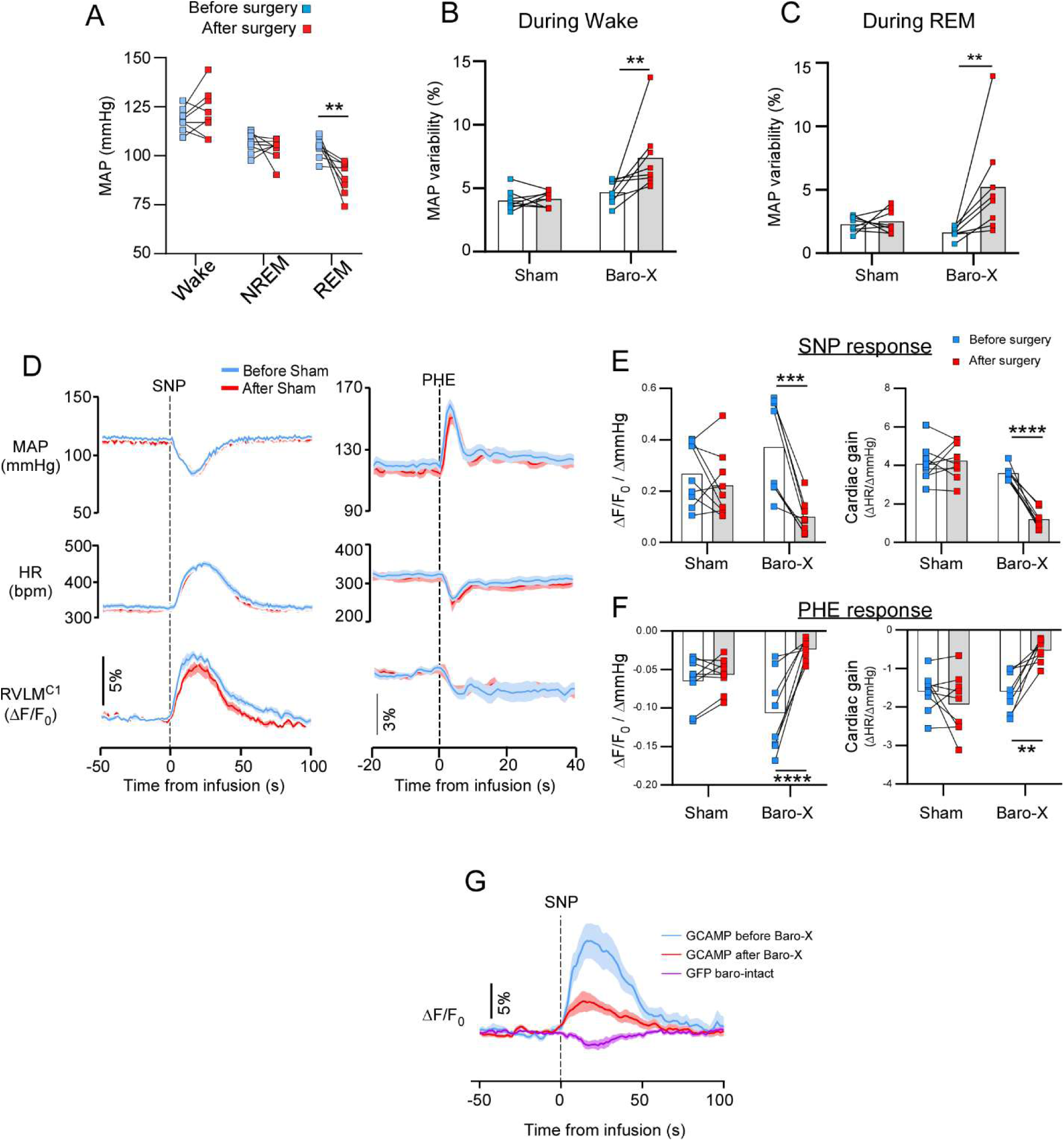
Effect of Sino-aortic denervation on RVLM^C1^ activity during BP perturbations and sleep-wake transitions. **A)** MAP in Baro-X rats during NREM sleep, wakefulness, and REM sleep. Two-way RM ANOVA, Time point (before vs. after surgery) × sleep-wake state, F (2, 14) = 6.26, P=0.012; sleep-wake state, F (1, 7) = 5.32, P=0.054; Time point, F (2, 14) = 47.43, P<0.0001. Asterisk indicate results of Šídák’s multiple comparisons test. ** p<0.01 **B.** MAP variability in Sham (N=8) and Baro-X (N=8) during stable wakefulness. Two-way RM ANOVA, Surgery type (sham vs. Baro-X) × recording session (Pre vs. post-surgery), F (1, 14) = 5.6, P=0.033; Surgery type, F (1, 14) = 12, p=0.003; recording session, F (1, 14) = 6.6, p=0.022. Asterisk indicate results of Šídák’s multiple comparisons test. ** p<0.01 **C.** MAP variability in Sham (N=8) and Baro-X (N=8) rats during stable REM sleep. Two-way RM ANOVA, Surgery type × recording session, F (1, 14) = 5.6, P=0.032; Surgery type, F (1, 14) = 1.9, p=0.193; recording session, F (1, 14) = 7.2, p=0.018. Asterisk indicate results of Šídák’s multiple comparisons test. ** p<0.01 **D.** MAP, HR and ΔF/F_0_ in response to SNP and PHE before and after Sham surgery (n=9). **E.** Grouped data for ΔF/F_0_ gain ((ΔF/F_0_)/ΔmmHg) and HR gain (ΔHR/ΔmmHg) in response to SNP for sham (n=9) and Baro-X (n=8) rats. For ΔF/F_0_ gain- RM two-way ANOVA, Surgery type × recording session, F (1, 15) = 11, p=0.004; Surgery type, F (1, 15) = 0.028, p=0.87; recording session, F (1, 15) = 23, p<0.001. For HR gain - RM two-way ANOVA, Surgery type × recording session, F (1, 15) = 42, p<0.001; Surgery type, F (1, 15) = 34, p<0.001; recording session, F (1, 15) = 31, p<0.001. Asterisk indicate results of Šídák’s multiple comparisons test- *** p<0.001, **** p<0.0001 **F.** Grouped data for ΔF/F_0_ gain and HR gain in response to PHE for sham (n=9) and Baro-X (n=8) rats. Mean ± SEM. For ΔF/F_0_ gain-RM two-way ANOVA, Surgery type (sham vs. Baro-X) × recording session (Pre vs. post), F (1, 15) = 20, p<0.001; Surgery type, F (1, 15) = 0.012, p=0.74; recording session, F (1, 15) = 30, p<0.001. For HR gain - RM two-way ANOVA, Surgery type × recording session, F (1, 15) = 17, p<0.001; Surgery type, F (1, 15) = 11, p=0.005; recording session, F (1, 15) = 4.7, p=0.046. Asterisk indicate results of Šídák’s multiple comparisons test- *** p<0.001, **** p<0.0001. **G.** ΔF/F_0_ in response to SNP before and after Baro-X surgery in GCAMP expressing rats (n=9) and in baro-intact rats expressing GFP (N=8) in RVLM^C1^ neurons. Mean ± SEM.

**Supplementary figure 4.**
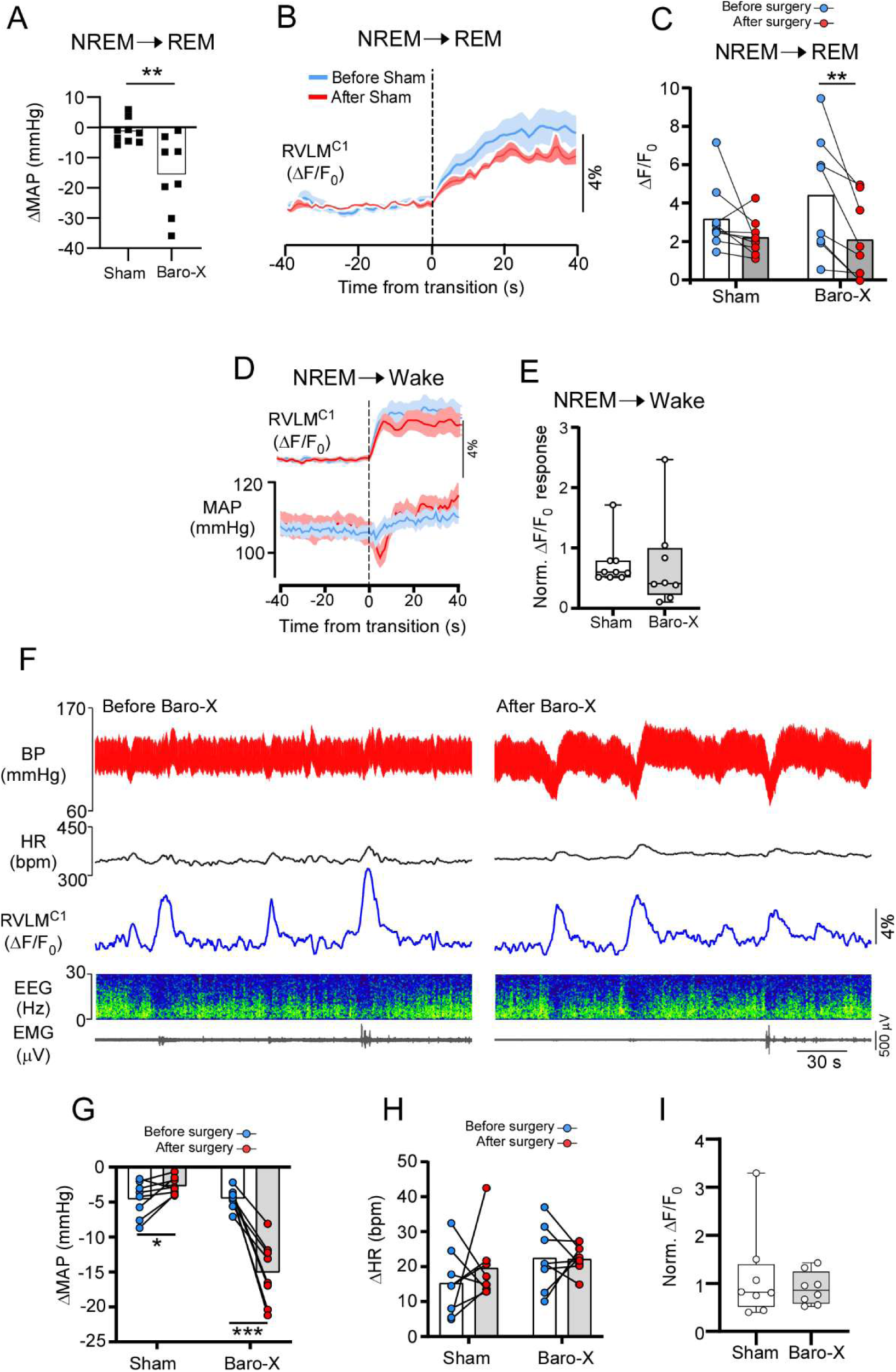
Effect of Sino-aortic denervation on RVLM^C1^ activity during sleep-state transitions and microarousals. **A.** Grouped data for the ΔMAP occurring during NREM-to-REM transitions in Sham (N=8) and Baro-X (N=8). Unpaired t test, t=3.184, df=15, P= 0.0062. **B.** Mean ΔF/F_0_ and MAP during NREM-to-REM transitions before and after Sham surgery (n=9). Mean ± SEM. **C.** Grouped data for ΔF/F_0_ during NREM-to-REM transitions for sham (n=9) and Baro-X (n=8) rats. RM two-way ANOVA, Surgery type (sham vs. Baro-X) × recording session (Pre vs. post), F (1, 15) = 2.4, p=0.14; Surgery type, F (1, 15) = 0.38, p=0.55; recording session, F (1, 15) = 14, p=0.002. Asterisk indicate results of Šídák’s multiple comparisons test-** p<0.01. **D.** Time course of ΔF/F_0_ and MAP during NREM-to-wake transitions before and after Baro-X surgery (n=8). Mean ± SEM. **E.** Normalized ΔF/F_0_ (ratio of post-to pre-surgery) for NREM-to-wake transitions in Sham (n=9) and Baro-X (n=8) cases. Mann Whitney test, U=35, p=0.97. **F.** Representative recordings of BP, HR, RVLM^C1^ activity (ΔF/F_0_), EEG, and EMG during microarousals before and after Baro-X in the same rat. **G.** Grouped data for ΔMAP during microarousal in Sham (N=8) and Baro-X (N=8) operated rats. RM two-way ANOVA, Surgery type × recording session, F (1, 14) = 49.33, p<0.0001; Surgery type, F (1, 14) = 33.75, p<0.0001; recording session, F (1, 14) = 23.97, p<0.0001. Asterisk indicate results of Šídák’s multiple comparisons test-* p<0.05, *** p<0.001. **H.** Grouped data for ΔHR during microarousal in Sham (N=8) and Baro-X (N=8) operated rats. RM two-way ANOVA, Surgery type × recording session, F (1, 14) = 0.49, p=0.5; Surgery type, F (1, 14) = 3.3, p=0.09; recording session, F (1, 14) = 0.39, p=0.55. **I.** Normalized peak ΔF/F_0_ (ratio of post-to pre-surgery) during microarousals in Sham (n=8) and Baro-X (n=8) cases. Mann Whitney test, U=31, p=0.96.

**Supplementary figure 5.**
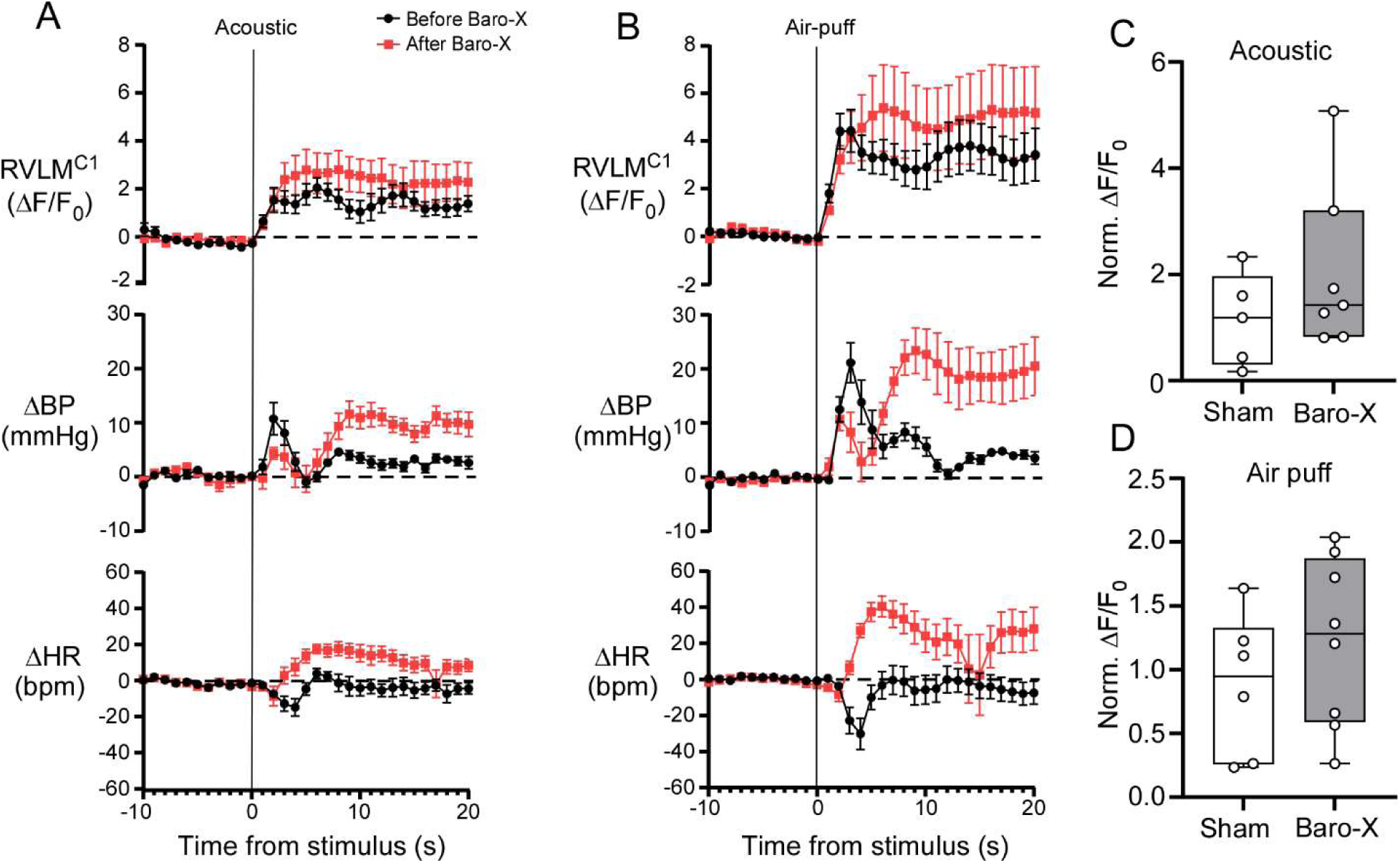
Effect of Sino-aortic denervation on RVLM^C1^ activity during acoustic stimulation and air-puff. **A.** Grouped data for ΔF/F_0_, ΔMAP, and ΔHR in response to arousal from NREM induced by acoustic stimulation in Baro-X (n=7) rats before and after surgery. Mean ± SEM. ΔMAP for acoustic stimulation-RM two-way ANOVA, time × recording session, F (4.4, 26) = 5.6, P<0.001; time, F (4.3, 26) = 13, P=0.001; recording session, F (1.0, 6.0) = 14, P=0.009. ΔHR for acoustic stimulation-RM two-way ANOVA, time × recording session, F (3.7, 22) = 5.9, P=0.003; time, F (2.6, 16) = 3.8, P=0.037; recording session, F (1.0, 6.0) = 13, P=0.011. **B.** Grouped data for ΔF/F0, ΔMAP, and ΔHR in response to arousal from NREM induced by air-puff in Baro-X (n=7) rats before and after surgery. Mean ± SEM. ΔMAP for air-puff-RM two-way ANOVA, time × recording session, F (1.6, 11) = 8.4, P=0.008; time, F (2.3, 16) = 17, P<0.001; recording session, F (1.0, 7.0) = 11, P=0.013. ΔHR for air-puff - RM two-way ANOVA, time × recording session, F (40, 280) = 8.9, P<0.001; time, F (40, 280) = 2.1, P<0.001; recording session, F (1, 7) = 20, P=0.003. **C.** Normalized peak ΔF/F0 (ratio of post-to pre-surgery) during acoustic stimulation in Sham (n=5) and Baro-X (n=7) cases. Mann Whitney test, U=11, p=0.34. **D.** Normalized peak ΔF/F0 (ratio of post-to pre-surgery) during air-puff in Sham (n=5) and Baro-X (n=7) cases. Mann Whitney test, U=18, p=0.83.

**Supplementary figure 6.**
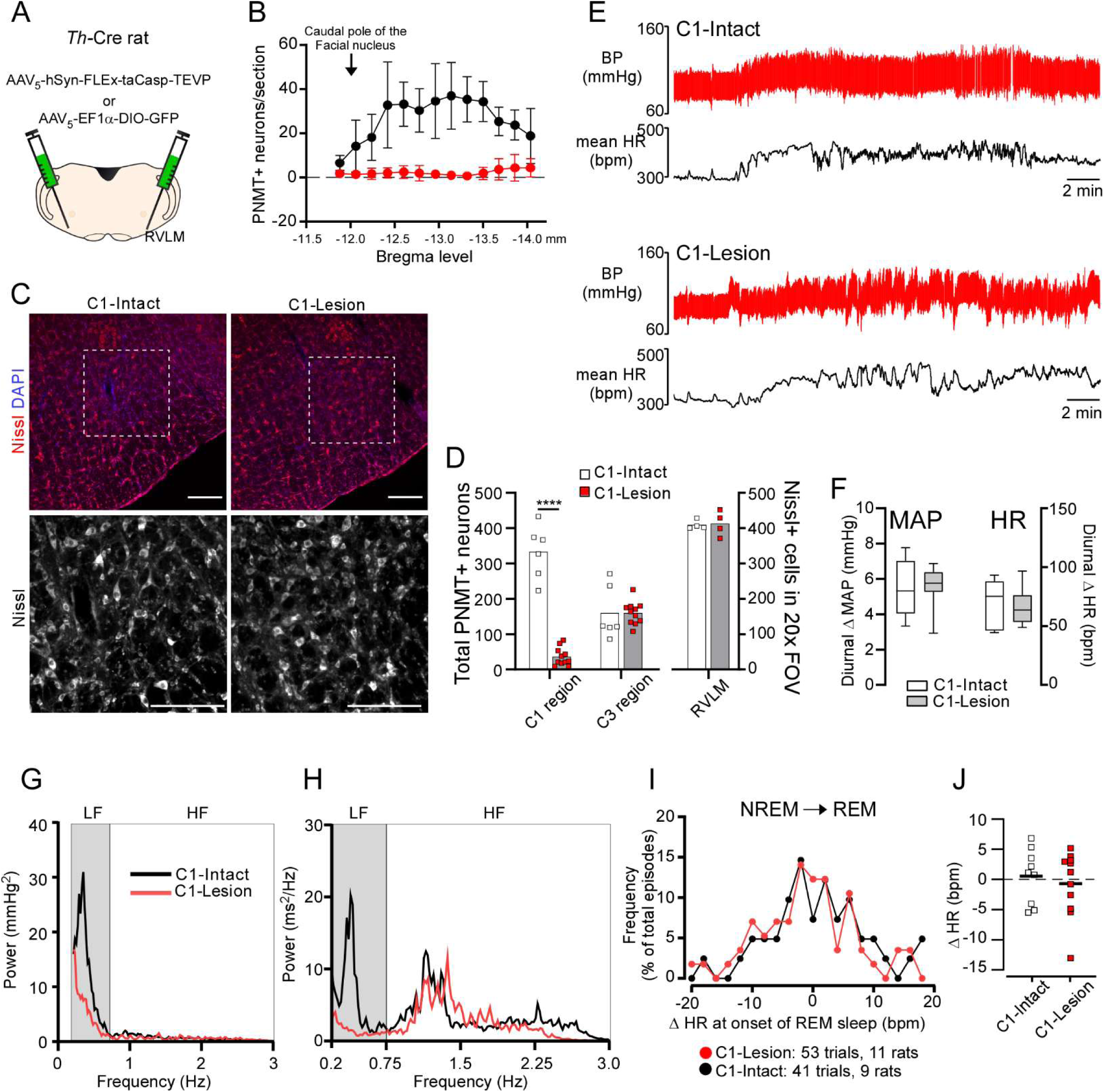
Effect of ablation of RVLM^C1^ activity on BP. **A.** Approach for genetically targeted ablation of RVLM^C1^ neurons. **B.** Rostral-caudal distribution of the TH+ neurons in the ventrolateral medulla in C1-Lesion (N=11) and C1-Sham (N=5) rats. **C.** Images of Nissl and DAPI staining in the RVLM at the site of virus injection. Scale bar: 100 µm **D.** Grouped data for counts of PNMT+ in the RVLM (C1 region) and in the dorsal medulla (C3 region) in C1-Sham (n=6) and C1-Lesion rats (n=11). C1 region-unpaired Student’s t-test, t=12.50, df=15, P<0.0001. C3 region-unpaired Student’s t-test, t=1.69, df=15, P= 0.11. Grouped counts of Nissl+ cell bodies in the RVLM in a 0.0625 mm^2^ field of view. Unpaired t-test, t=0.1996, df=6, p=0.85. **E.** Representative recordings of BP and HR in a C1-Sham (upper panel) and C1-Lesion (lower panel) rat. **F.** Grouped data for ΔMAP and ΔHR between mid-light and mid-dark phase. For MAP, Unpaired t-test, t=0.1081, df=13, p=0.92. For HR, Unpaired t-test, t=0.5041, df=13, p=0.63. **G.** Representative spectra of heart rate variability in during wakefulness in a C1-Sham and C1-Lesion rat. **H.** Representative spectra of systolic BP in during wakefulness in a C1-Sham and C1-Lesion rat. **I.** Frequency distribution of ΔHR across NREM-to-REM transitions in C1-Intact and C1-Lesion cases. **J.** Grouped data for ΔHR during NREM-to-REM transitions in C1-Intact (n=9) and C1-Lesion (n=11) cases. Unpaired t-test, t=0.5588, df=18, p=0.58.

**Supplementary Table 1.**

|  | Male |  | Female |  |
| --- | --- | --- | --- | --- |
|  | **Control** | **C1-Lesion** | **Control** | **C1-Lesion** |
| **Parameter (Mean ± SD)** | **n=6** | **n=8** | **n=3** | **n=3** |
| MAP mmHg | 110 ± 10 | 109 ± 7 | 110 ± 7 | 100 ± 14 |
| HR bpm | 333 ± 24 | 331 ± 23 | 365 ± 23 | 364 ± 6 |
| MAP Variability (%) | 4.6 ± 0.3 | 5.4 ± 1 | 4.7 ± 0.3 | 5.1 ± 1 |
| LF-SAP mmHg<sup>2</sup> | 5.2 ± 2 | 2.1 ± 1 | 5.3 ± 3 | 2.4 ± 1 |

## Methods

### Subjects

All experiments were conducted in accordance with the National Institutes of Health’s Guide for the care and use of laboratory rats and approved by the University of Virginia Animal Care and Use Committee (protocol #4312). We used 46 adult male and female transgenic Sprague Dawley rats that express Cre-recombinase under the control of the endogenous tyrosine hydroxylase (*Th*) promoter, ie., TH-Cre. This model possesses a targeted insertion of (IRES)-cre immediately after the translational stop in the open reading frame of *Th* (HsdSage:SD-TH^em1(IRES-Cre)Sage^, RRID: RGD_12905029, originally purchased from Sage, now available from Inotiv). The selectivity of the Cre expression in C1 neurons in TH neurons has been validated in our previous studies (Souza et al., 2019, 2020, 2022b) as well as the present study. Rats weighed 250-300 g at the time of virus microinjections and 300-550 g at the time of experiments. Rats were housed at 23°C-24°C under a standard artificial 12-hour light-dark cycle with water and food provided *ad libitum*.

### Viral vectors

For the fiber photometry experiments, we used a Cre-dependent vector encoding GCaMP7s (Dana et al., 2019) (AAV_1_-hSyn-FLEx-jGCaMP7s-WPRE, Addgene plasmid: 104491, purchased from UPenn Vector Core, virus titer for injection: 4.7 × 10^11^ GC/mL). To induced cell-autonomous apoptosis of RVLM^C1^ neurons, we used AAV_5_-EF1α-FLEx-taCasp3-TEVp (Yang et al., 2013)(Addgene plasmid: 45580, purchased from UNC Vector Core, virus titer for injection: 4.6 × 10^12^ vg/mL). We used a Cre-dependent virus encoding eYFP as a control virus for GCAMP and lesion experiments (AAV_2_-EF1α-DIO-eYFP, purchased from UNC Vector Core, titer injected: 4.0 × 10^12^ vg/mL).

### Surgical preparation

All surgical procedures were conducted under aseptic conditions, and body temperature was maintained at 37°C with a servo-controlled heating pad. The depth of anesthesia was judged by the lack of withdrawal reflex following a firm tail pinch and the absence of a corneal reflex. For brain injections and electroencephalogram and neck electromyogram (EEG/EMG) electrode implantation, rats were anesthetized with a mixture of ketamine (75 mg/kg), xylazine (5 mg/kg), and acepromazine (1 mg/kg, i.m.). Additional anesthetic was administered if required (25% of the original dose, i.m.). Brain injections were performed on a stereotaxic frame (David Kopf Instruments), with the bite bar set at 3.5 mm below the interaural line for a flat skull. In all rats, incisions were closed in two layers using absorbable sutures for internal closures, and steel clips and VetClose tissue adhesive for skin. An analgesic (ketoprofen, 3-5 mg/kg, s.c.) was administered on the day of surgery and every 24-hour for 3 days following surgery. All experiments were performed after at least 1 month of recovery from virus microinjections.

### Virus microinjections, optical fiber placement, and electroencephalogram (EEG) and neck electromyogram (EMG) head stage implantation

RVLM microinjections were performed with electrophysiological mapping of the facial motor nucleus (Brown and Guyenet, 1985). The microinjections were performed using a glass micropipette and a picopump (Picospritzer, Parker Hannifin). For RVLM^C1^ neuron targeting with GCaMP7s, one microinjection (60 nl; left side of the brain) were placed 1,800 µm lateral to the midline, 100 µm below and 400 µm caudal from the caudo-ventral pole of the facial nucleus.

During the same surgery a custom-made EEG and neck EMG electrodes was implanted. The head stage consisted of a 6-pin plastic headstage (P1 Technologies) fitted with 5 amphenol pins connected to 5 Teflon-insulated multistrand stainless-steel wires (coated diameter 0.009-inch, A-M Systems); three wires were soldered to stainless steel jeweler screws (0-86, 3 mm shaft) and the remaining two wires were stripped of Teflon at the tips (2 mm of bare wire). During surgery, stainless-steel jeweler screws were implanted in the left and right parietal bone (6 mm caudal of bregma, 3.5 mm lateral of midline) and the right frontal bone (3 mm rostral of bregma, 3 mm lateral of midline) for EEG and the two stripped bare wires were implanted in the dorsal layer of the neck muscle and secured with 4-0 silk sutures for EMG. After this procedure a low-fluorescence optic fiber cannula (400 µm core, 0.57 NA, Doric Lenses) was implanted with the tip placed 300-400 µm dorsal to the virus injection site and then the head stage and fiber optic cannula were secured to the skull with dental cement.

For lesion experiments, AAV_5_-EF1α-FLEx-taCasp3-TEVp or AAV_2_-EF1α-DIO-eYFP was injected in the RVLM in 3 sites separated by 400 µm along the rostro-caudal axis, with the most rostral site placed 1800 µm lateral to the midline, 100 µm below and 400 µm caudal from the caudo-ventral pole of the facial nucleus. Rats where then fitted during the same surgery with an EEG/EMG head stage as described above.

### Blood pressure measurements and venous catheter implantation

Four to five weeks after virus microinjections and fiber implantation, rats were re-anesthetized with 2.5% isoflurane in pure oxygen. A small incision was made on the right leg for the implantation of a polyethylene catheter (PE-10 connected to a PE-50, Clay Adams) in the femoral vein for drug infusion. The catheter was tunneled under the skin and exteriorized in the dorsal aspect of the neck. On the same leg, the femoral artery was exposed and a radiotelemetry probe (PA-C10, Data Sciences International) was implanted to monitor arterial pressure.

### Fiber photometry

Fiber photometry was performed as described previously (Souza et al., 2022). We used a 470 nm LED modulated at 211 Hz for GCaMP7s excitation and a 405 nm LED modulated at 531 Hz as a control fluorescent channel to corrected for photobleaching and movement artifacts.

Modulation and demodulation of LEDs and the emitted fluorescence was performed using two lock-in amplifiers (SR810 DSP, Stanford Research Systems). Both light sources were coupled through a fluorescent minicube (Doric Lenses) into a low-fluorescence optic fiber cable (400 µm core, 0.57 NA, Doric Lenses) multiple-mode fiber optic (Doric Lenses, 400 µm, 0.57 NA). The mean power output of each LED was set at ∼12 µW when measured at the tip of the fiber connected to the rat using a power meter (Thorlabs). Fluorescence signals were collected by a photoreceiver (Newport) before demodulation and acquisition. Prior to experiments, recording quality was assessed by intravenous injection of sodium nitroprusside (SNP in saline, 5 µg·kg^-1^ in a 0.1mL bolus, Millipore-Sigma, product # 228710) to unload the arterial baroreceptors; an increase in fluorescence of less than 2% from baseline following SNP was judged insufficient and rats were excluded from further experiments.

### Sino-aortic denervation

After the baseline recordings, rats (n=8, 5 male and 3 female) rats were anesthetized with a mixture of ketamine (75 mg/kg), xylazine (5 mg/kg), and acepromazine (1 mg/kg, i.m.) to perform sino-aortic denervation as previously described (Krieger, 1964). A ventral mid-line incision on the neck was followed by retraction of glandular tissue, connective tissue and muscle to expose the superior laryngeal nerve, superior cervical ganglion, vagus nerve and carotid bifurcation. The carotid sinuses were denervated by stripping of nerve fibers from the carotid bifurcation. Wounds were closed with absorbable sutures and surgical clips. An analgesic (ketoprofen, 3-5 mg/kg, s.c.) was administered on the day of surgery and every 24 hour for 3 days following surgery. Recordings experiments were performed 7 days after the surgery. Efficacy of the sino-aortic denervation was established by the absence of fall in heart rate (HR) during an increase in BP produced by intravenous phenylephrine hydrochloride (PE in saline, 5 µg·kg^-1^ in a 0.1mL bolus, Millipore-Sigma, product # 1533002). Other parameters, such as hypotension during REM sleep, increased BP lability and reduced reflex tachycardia in response to hypotension also confirmed the efficacy of the denervation (Krieger, 1964; Silveira et al., 2008; Amorim et al., 2016).

### Protocol for fiber photometry recordings in plethysmograph chambers

Experiments conducted in plethysmography chambers were performed between 0900 and 1600 at an ambient room temperature of 27°C-28°C. Rats were habituated to the experimental conditions for 2 days before recording as described previously. On the day of the recording, rats were briefly anesthetized with isoflurane (2.0% in room air for 3 min) to connect the fiber optic implant to connecting optical fiber, the EEG/EMG head stage to an amplifier, and the venous catheter to catheter connected to a syringe containing sterile saline in the case of fiber photometry recordings, and the EEG/EMG head stage to an amplifier in the case of lesion experiments. Rats were then placed in an unrestrained whole-body plethysmography chamber (5 L volume, EMKA Technologies). Bias flow (2.5 L/min) was maintained using mass flow controllers (Alicat Scientific). Experiments were performed in room air conditions (F_I_O_2_ = 0.21, F_I_CO_2_ = 0.0, balance N_2_). Rats were allowed at least 1 hour before recordings were initiated. Recording sessions lasted up to 6 hours.

First, the arterial baroreflex was evaluated by intravenous infusion of SNP and PHE (5-10 µg·kg^-1^ in a 0.1-0.2 mL bolus). This was performed up to three times per drug. After this, physiological variables were recorded for 4-5 hours while the rats were left undisturbed in the chamber, in which they spent the majority of the time sleeping. Our previous data shows that time spent in wakefulness, NREM and REM sleep in these conditions is comparable to recordings in a home-cage environment ((Souza et al., 2019) compared with (Qiu et al., 2010)). After sufficient data was collected for analysis of sleep-wake transitions, we induced arousal using acoustic stimulation and air-puff delivered during NREM sleep 2 to 3 times for each stimulus modality. For acoustic stimulation, a mini speaker was placed inside of the recording chamber underneath the rat. The speaker was driven by Spike 2 software using a 250 Hz sinusoidal wave for 2 s at 100 dBA. For the air-puff, a cannula connected to compressed air canister was inserted through a side-port in the chamber to deliver a short air-puff lasting <1 s against the torso of the sleeping rat.

### Protocol for characterization of cardiovascular phenotype after RVLM^C1^ lesion

The effect of RVLM^C1^ lesion on cardiovascular function was examined in three different recording sessions. First, we characterized BP and heart rate with rats individually housed in cages (dimensions: 36 x 26 x 18 cm) with *ad libitum* access to food and water under a standard artificial 12-hour light-dark cycle at 23°C-24°C. Data was collected continuously for 3 days with grouped data representing the final day of recording. Next, we characterized BP and heart rate with recordings of EEG/EMG recordings inside a plethysmography chamber to identify periods of sleep and wake and to assess the cardiovascular response to acoustic and air puff stimulation as described above. Third, we characterized BP and heart rate parameters in a large custom-made chamber (60 cm × 30 cm × 30 cm) with a 12-hour light-dark cycle and food and water provided *ad libitum*. The chamber was divided into two spaces by a 15 cm high partition that rats would jump over during exploration. Recording sessions were conducted between 1800 and went until 1200 (lights off 2000) with behavioral activity recorded using an infrared (night-vision) camera.

### Data acquisition

All signals were digitized with a CED 1401 A/D acquisition system using Spike 2 (Version 9, Cambridge Electronic Design, UK). The analog readout for the demodulated 470 nm and 405 nm channels were acquired at 1 kHz and smoothed with a time constant of 1 s. EEG and EMG signals were bandpass filtered (EEG: 0.1-100 Hz, EMG: 100-300 Hz) and amplified (×2,000) and then digitized at 1 kHz. Radiotelemetry signals were first converted from a digital to analog signal (R11CPA, DSI) incorporating a reference to ambient pressure (APR1, DSI). The analog output was then digitized and acquired at 1 kHz using a CED 1401 A/D acquisition system.

Heart rate was generated from the pulsatile BP signal using the peak find function in the Spike 2 software.

### Analysis

#### Blood pressure, heart rate and variability index

Mean arterial pressure (MAP) and heart rate (HR) were derived from pulsatile BP signals. We used the coefficient of variation (CV) as an index of variability of MAP, HR and fluorescence. CV was calculated by the formula: CV = SD of the data set/mean of the data set × 100%, and expressed as a percentage. For 2-hour data we extract average SD for each 2-hour segment of mean BP and applied the formula. For all other MAP variability, MAP was extracted at 1 Hz and the SD of this data was used to determine CV.

#### Fiber photometry

ΔF/F_0_ was derived from the demodulated emission of the 470 nm and 405 nm channel using the following formula: ΔF/F_0_ = (ΔF_470_/F_0_)/(ΔF_405_/F_0_) × 100. ΔF/F_0_ was extracted from Spike2 at 1 Hz and plotted as mean ± S.E.M. Each data point in the grouped data reflects a within-animal average of 2-5 trials for each condition or stimulus. In some cases, ΔF/F_0_ is presented as a z-score [z= (value – mean)/SD]. To evaluate the effects of sino-aortic denervation on RVLM^C1^ activity, we generated a ratio of pre-surgery ΔF/F_0_ normalized to post-surgery ΔF/F_0_ for each stimulus or condition for the sham and Baro-X groups. This ratio accounts for differences in the recording quality across sessions and cell transduction while retaining quantitative values for ΔF/F_0_ statistical comparisons between groups. Unnormalized values of ΔF/F_0_ pre- and post-surgery are presented in the supplementary materials.

#### Definition of sleep-wake stages

Sleep-wake state was characterized is either NREM, REM or wake based on EEG/EMG recordings according to conventional criteria. Non-rapid eye movement (NREM) sleep was identified by slow-wave EEG activity (delta, 1–4 Hz) and low EMG activity. Rapid eye movement (REM) sleep was defined by a stable theta rhythm (6–8 Hz) and EMG atonia. Wakefulness/arousal was characterized by the absence of slow-wave EEG activity, generalized EEG desynchronization, heightened and variable EMG activity.

#### Baroreflex gain

The gain of RVLM^C1^ activity and heart rate during BP perturbations by SNP and PHE were calculated as the peak or nadir changes in fluorescence ((ΔF/F_0_)/mmHg)) and heart rate (Δbpm/mmHg) following intravenous infusion of SNP or PHE. A baroreflex gain of HR of 1bpm/mmHg or less was used as an inclusion criterion for rats in the Baro-X group; one rat not meeting this criterion was excluded from the grouped data. All rats in the underwent sham surgery group were included in the grouped data.

#### Spectral analysis of systolic arterial pressure (SAP) and heart rate (HR)

The spectral analysis of systolic arterial pressure (SAP) and pulse interval (PI) was performed to quantify the sympathetic modulation of BP and HR (Pagani et al., 1986; Malliani et al., 1991). Data were analyzed using the software (CardioSeries v2.7; https://www.danielpenteado.com). SAP and PI signals were derived from segments of BP for each sleep-wake stage. Spectral analysis was used to evaluate variability in the frequency domain where power spectral density was estimated using the fast Fourier transform, and the resulting spectra were integrated over two frequency bands: low frequency (LF, 0.2–0.75 Hz) and high frequency (HF, 0.75–3.0 Hz). LF component of SAP and HR is indicative of sympathetic modulation of blood pressure and heart, respectively and the HF component of HR is an indicative of parasympathetic modulation of the heart. The ratio of LF to HF (LF/HF) served as an indicator of sympathovagal balance.

#### Video analysis of behavior

We focused on three stereotyped behaviors that were easily characterized in our video recordings; rearing, jumping over the barrier, and grooming. Rearing events consisted of the rat raising up to stand on its hind paws in a single movement with or without contacting the walls of the chamber with their fore paws. A jumping event consisted of the rat crossing the partition without pause. For rearing and jumping, the onset of the behavior shown in the grouped data refers to the time point at which the front paws of the rat leave the ground. Grooming consisted of the rat turns its body to groom the fur on its back for at least 10 s, the onset of this behavior refers to the to the time point at which the rat initiates a sequence of grooming. Changes in BP and HR were analyzed in at least three trials of each behavior.

#### Histology

At the end of the experiment, rats were deeply anesthetized with a mixture of ketamine (112.5 mg/kg), xylazine (7.5 mg/kg), and acepromazine (1.5 mg/kg, i.m.) and then transcardially perfused first with 300 mL of heparinized saline and then with 300 mL of 10 % formalin. Brains were removed and postfixed in the same fixative for 12-16 hours at 4°C. Brains were then sectioned the next day on a vibratome (VT-1000S, Leica Biosystems) in the transverse plane at 30 μm, and brain slices were collected in 6 series and then stored in cryoprotectant at −20°C. Immunostaining procedures were performed on free-floating sections at room temperature unless noted otherwise.

A 1-in-6 series of sections were rinsed, then blocked in a solution containing 100 mM tris, 150 mM saline, 10% horse serum (v/v) 0.1% Triton-X (v/v), then incubated with primary antibodies for 60 min at room temperature then 4°C overnight. The next day, sections were rinsed and then incubated with secondary antibodies for 60 min and rinsed again before mounting on slides. Slides were cover slipped with ProLong Gold anti-fade mounting medium with DAPI (P36931, Thermo Fisher Scientific). Primary antibodies: Sheep anti-TH (1:1k, Millipore Sigma, catalog #AB1542, RRID: AB_90755), Rabbit anti-PNMT (1:5k, Gift from Dr. M. Bohn, RRID:AB_2315181), Mouse anti-TH (1:10k, Millipore Sigma, catalog T1299, RRID:AB_477560). Secondary antibodies were obtained from Jackson ImmunoResearch Laboratories: AlexaFluor-488 AffiniPure Donkey Anti-Chicken IgY (IgG) (H+L)(catalog #703-545-155, RRID:AB_2340375), Cy3 AffiniPure F(ab’)2 Fragment Donkey Anti-Mouse IgG (H+L)(catalog #715-166-150, RRID:AB_2340816), Cy3 AffiniPure Donkey Anti-Rabbit IgG (H+L) (catalog #711-165-152; RRID: AB_2307443)AlexaFluor-488 AffiniPure F(ab’)2 Fragment Donkey Anti-Mouse IgG (H+L)(catalog #715-546-150, RRID:AB_ 2340849). All secondary antibodies were used at a dilution of 1:500. Nissl staining using Neurotrace (Invitrogen, catalog # N21482, RRID: AB_2620170) was performed as per manufacturer’s instructions, with all steps conducted at room temperature. In brief, tissue sections were mounted and dried on a slide, then rehydrated in phosphate-buffered saline (PBS) for 40 minutes. Following rehydration, the tissue was then washed in PBS plus 0.1% Triton X-100 (PBS-T), followed by two brief washes in PBS. The tissue was then incubated in a 1:200 dilution of NeuroTrace 530 for 20 minutes. Tissue was then rinsed in PBS-T for 10 minutes and then PBS twice briefly and finally in PBS for 2 hours.

#### Neuronal mapping

Neuronal mapping was conducted using a motor driven stage with Neurolucida software (version 2021, MBF Bioscience) with an AxioImager M2 microscope (Carl Zeiss). Digital photomicrographs were acquired with the same microscope in grayscale using a C11440 Orca-Flash 4.0LT digital camera (Hamamatsu). The selectivity of GCAMP7s expression in RVLM^C1^ neurons was evaluated by the colocalization of native GCAMP7s fluorescence with TH. The extent of cell loss of RVLM^C1^ neurons and C3 neurons in the dorsomedial medulla was determined by blinded counting of PNMT^+^ cell profiles in the ventral medulla for C1 and dorsal medulla for C3 region between −11.5 mm and −14.0 mm caudal of bregma. The selectivity of C1 cell loss was also evaluated by counting Nissl^+^ neuronal profiles in the ventrolateral medulla by a blinded observer. Only cell profiles that included a nucleus in the plane of section were counted and/or mapped.

#### Statistics

Statistical comparisons were performed using Prism software (version 10, GraphPad). Following tests for normality (D’Agostino-Pearson or Shapiro-Wilk), significant differences were determined using repeated measures one-way and two-way ANOVA with Tukey’s or Dunnet’s post-test and unpaired Student’s t-test as indicated in figure legends. Correlations were evaluated using Pearson correlation coefficient. All F and p values for the interaction effects and individual treatment effects are reported in the figure legends. Data are reported as mean and range unless otherwise noted and differences were considered significant when p < 0.05.

## Author contributions

G.M.P.R.S. and S.B.G.A. designed research; G.M.P.R.S., H.T., D.S.S., and S.B.G.A. performed research; G.M.P.R.S., H.T, F.E.B, U.M.A, D.S.S., and S.B.G.A. analyzed data; G.M.P.R.S. and S.B.G.A. wrote the paper.

This work was supported by the National Institutes of Health grant HL148004 to S.B.G.A. The authors declare no competing financial interests.

